# The integrated stress response remodels the microtubule organizing center to clear unfolded proteins following proteotoxic stress

**DOI:** 10.1101/2022.02.13.480280

**Authors:** Brian Hurwitz, Nicola Guzzi, Anita Gola, Vincent F. Fiore, Ataman Sendoel, Maria Nikolova, Douglas Barrows, Thomas S. Carroll, H. Amalia Pasolli, Elaine Fuchs

## Abstract

When cells encounter stressful situations, they activate the integrated stress response (ISR), which limits total protein synthesis and redirects translation to proteins that help the cells to cope. The ISR has also been implicated in cancers, but redundancies in the stress-sensing kinases that trigger the ISR have posed hurdles to dissecting physiological relevance. To overcome this challenge, we targeted the regulatory node of these kinases, namely the S51 phosphorylation site of eukaryotic translation initiation factor eIF2α and genetically replaced eIF2α with eIF2α-S51A in squamous cell carcinoma (SCC) stem cells. While inconsequential under normal growth conditions, the vulnerability of this ISR-null state was unveiled when SCC stem cells experienced proteotoxic stress. Seeking mechanistic insights into the protective roles of the ISR, we combined ribosome profiling and functional approaches to identify and probe the functional importance of translational differences between ISR-competent and ISR-null SCC stem cells when exposed to proteotoxic stress. In doing so, we learned that the ISR redirects translation to centrosomal proteins that orchestrate the microtubule dynamics needed to efficiently concentrate unfolded proteins at the microtubule organizing center so that they can be cleared by the perinuclear degradation machinery. Thus, rather than merely maintaining survival during stress, the ISR also functions in promoting cellular recovery once the stress has subsided. This finding exposes a vulnerability to SCC stem cells that could be exploited therapeutically.

## Introduction

Eukaryotic cells rely on highly conserved pathways to dynamically control translational machinery in response to various stimuli. One such pathway, the integrated stress response (ISR), is triggered when one of four stress-sensing kinases becomes active: (1) Heme-regulated inhibitor kinase (HRI) activated by oxidative stress, such as arsenite; (2) Protein Kinase R (PKR) is activated by viral infections; (3) PKR-like endoplasmic reticulum kinase (PERK) is induced when unfolded proteins accumulate (proteotoxic stress) and when the endoplasmic reticulum (ER) becomes stressed; and (4) General control non-depressible 2 (GCN2) is induced in poor nutrient conditions, particularly amino acid deprivation (Costa-Mattioli & Walter, 2020; El-Naggar & Sorensen, 2018). Activation of any of these four kinases results in phosphorylation of serine 51 of eIF2α, a core component of the canonical translational initiation complex. Since phosphorated eIF2α blocks the guanine nucleotide exchange factor eIF2B from stimulating the eIF2-GTP-methionyl-initiator tRNA ternary complex, the translation of housekeeping mRNAs is decelerated, and initiation of new protein synthesis is dampened. Concomitantly, non-canonical translational mechanisms emerge to synthesize key stress response proteins that help restore physiological balance to the cell and aid in survival (Figure 1A) (Pakos-Zebrucka et al., 2016).

**Figure 1:**
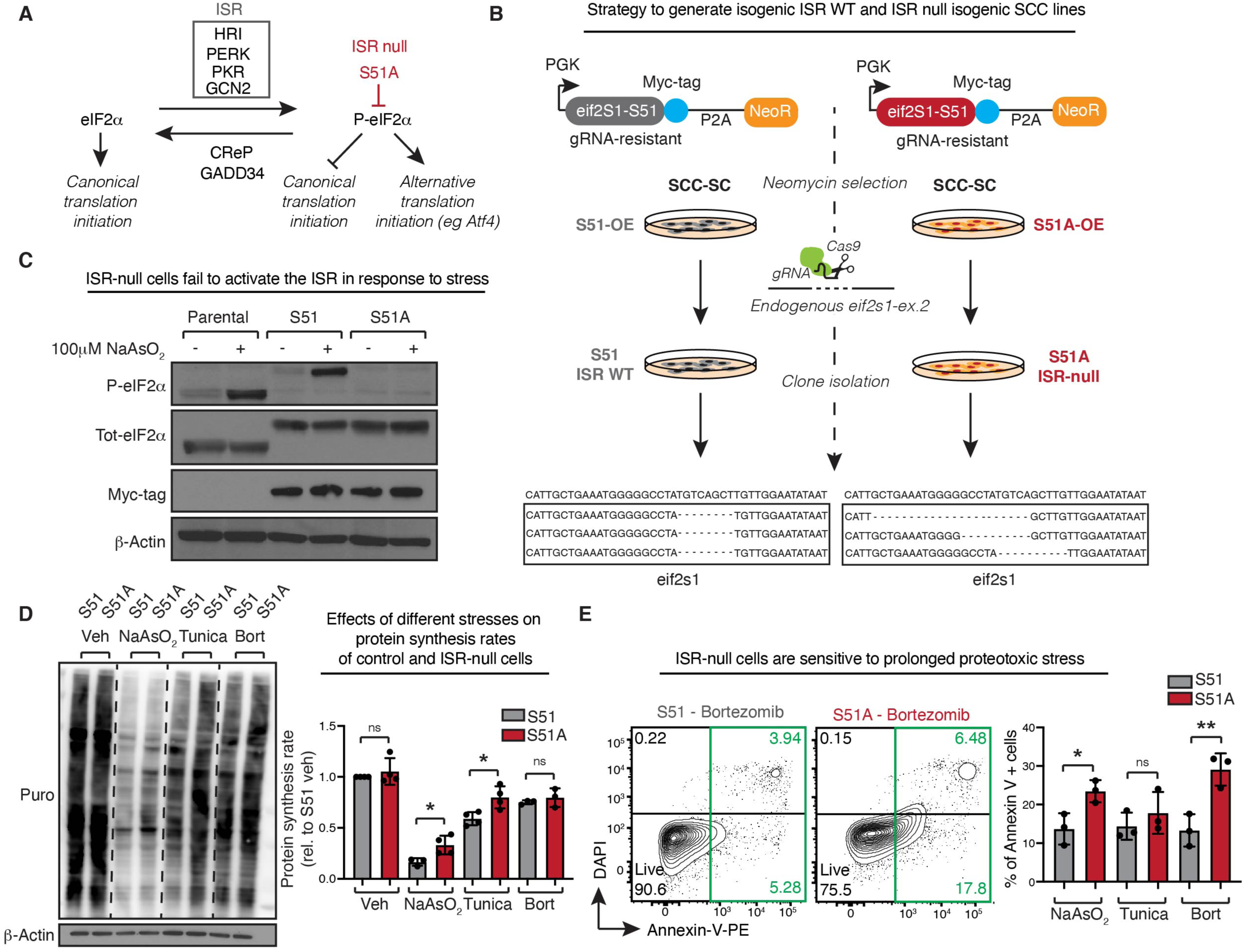
Generation of isogenic ISR-null SCC cell lines. A. Schematic of the integrated stress response pathway showing activating kinases, inhibitory phosphatases, and downstream effects. Mutation of eIF2α serine 51 to alanine renders cells insensitive to the ISR activating kinases (“ISR-null”). B. Reconstitution and knock out strategy to generate ISR-null SCC cell lines. Constructs encoding a myc-epitope tagged and CRISPR-Cas9 resistant eIF2α (either S51 or S51A) are packaged into lentivirus and integrated into a parental SCC stem cell (SCC-SC) line. Following neomycin selection, CRISPR-Cas9 is delivered by RNP transfection to selectively knock out endogenous eIF2α alleles. Clones grown from single cells are analyzed for knock out by deep sequencing the gRNA target locus. C. Representative immunoimmunoblot confirms both the replacement of the endogenous eIF2α with myc-tagged eIF2α and the failure of cells expressing eIF2α-S51A to activate the ISR. Cells were treated with 100 μM sodium arsenite for 6 hrs to induce oxidative stress. D. ISR-null cells show reduced capacity to activate the ISR as judged by global protein synthesis rate. ISR-WT and ISR-null cells are treated with 50 μM sodium arsenite (to induce oxidative stress), 100 ng/mL tunicamycin (to induce ER stress) or 100 nM bortezomib (to induce proteotoxic stress) for 6 hrs followed by incubation with puromycin at 1 μM for 30 minutes and anti-puromycin immunoimmunoblot. Bar graph on the right shows rate of protein synthesis ± standard deviation (SD) normalized to *β*-actin and relative to vehicle-treated S51 cells. Quantification of at least 3 independent biological replicates. * P<0.05 ns, no statistical significance (t test). E. ISR-null cells are more sensitive to stress. Early signs of apoptosis are measured by annexin V binding in control and ISR-null cells treated with 5 μM sodium arsenite, 100 ng/mL tunicamycin or 4 nM bortezomib for 24 hrs. Bar graph shows percentage of annexin-V positive cells ± SD of N=3 independent biological experiments. *P<0.05, **P<0.01, ns, no statistical significance (t test).

This paradoxical translational upregulation of a subset of mRNAs can be driven by several mechanisms including (1) lowering the dependency on the cap-binding complex eIF4F (Schuster & Hsieh, 2019), (2) initiating non-canonical translation within 5’ untranslated regions (5’ UTRs) that harbor dynamic N6-methyladenosine (m6A) RNA modification sites and/or 5’ upstream open reading frames (uORF) (Sendoel et al., 2017; Starck et al., 2016; Zhou et al., 2018) and finally (3) directly recruiting translation factors and ribosomes to the 5’ UTR of stress-responsive mRNAs (Guan et al., 2017). This selective repurposing of the translational machinery forms the foundation of a cellular stress response aimed at restoring homeostasis.

The best studied translational target of the ISR is *Atf4*, encoding activating transcription factor 4 (ATF4) whose translation is repressed in homeostatic conditions by the presence of inhibitory uORFs in its 5’ UTR. Upon stress, *Atf4* translation is unleashed, and the newly synthesized protein mediates transcriptional upregulation of stress response genes, thus acting as a crucial regulator in the balance of cell death and survival. In fact, known ATF4 targets include both cytoprotective and apoptotic factors. That a single ISR target can have such diverging and context-specific roles highlights the complexity of the cellular stress response and indicates the possibility that the ISR might translationally regulate additional cellular programs to coordinate survival, adaptation and recovery from stress.

At the interfaces of cellular proliferation, apoptosis, survival, and protein synthesis, the ISR has naturally emerged as an area of interest in cancer research. Although studies on the role of the ISR in cancer have been conflicting, the notion that the ISR is protective for cancer cells has gained traction in recent years (Ghaddar et al., 2021; Koromilas, 2015). This is in line with the observation that cancer cells experience hostile microenvironments characterized by nutrient deprivation and low oxygen availability. Additionally, the proliferative stress and elevated metabolic demands of cancer cells create an increased reliance on the cellular mechanisms that maintain proteostasis, a central function of the ISR (Cubillos-Ruiz et al., 2017). Indeed, proteasome inhibition as an anticancer therapy, is aimed at exploiting this vulnerability. That said, although proteasome inhibitors have become the standard of care for multiple myeloma and mantle-cell lymphoma (Manasanch & Orlowski, 2017), this strategy has been less effective in solid tumors, suggesting that these tumors may utilize mechanisms that enable them to cope, survive and recover in the face of proteotoxic stress, thereby evading therapeutics (Tian et al., 2021).

In the current study, we focus on the most common and life-threatening solid tumors, squamous cell carcinomas (SCCs), which affect stratified squamous epithelia of the skin, esophagus, lung and head and neck. To investigate the roles of the ISR in this cancer, we took advantage of our ability to culture the tumor-initiating stem cells from mouse skin SCCs and generated primary, clonal cell lines in which the endogenous eIF2α alleles were replaced by myc-epitope tagged but otherwise fully functional versions of either wild-type eIF2α or eIF2α-S51A (ISR-null). Although unable to mount an ISR in the face of stress, ISR-null SCC stem cells formed tumors that were similar in morphology and proliferation characteristics to controls. However, both *in vivo* and *in vitro,* when challenged with proteotoxic stress, ISR-null SCC cells fared considerably more poorly than their ISR-competent counterparts.

While the HRI and PERK kinases have emerged as major players in sensing misfolded proteins (Abdel-Nour et al., 2019; Harding et al., 1999), a detailed understanding of how the ISR promotes proteostasis is lacking. To probe into the mechanisms that link the ISR to proteostasis, we coupled genetics, cell biology, pharmacological inhibitors, ribosomal profiling and finally functional analyses. We traced the connection to a group of centrosomal proteins that become selectively translated in response to prototeoxic stress and which act by strengthening the organizing center for the microtubule dynamics that are needed to efficiently amass unfolded proteins at the pericentrosomal locale, where they can be efficiently targeted for destruction and clearance. Our findings add a new dimension—microtubule dynamics-- to the role of the ISR not only in stress, but also in the recovery of cells to stress. In so doing, our findings also expose a hitherto unappreciated vulnerability of cancer cells when they are unable to mount an ISR in the face of proteotoxic stress.

## Results

### Generation of ISR-null SCC cells

In order to directly test the role of the integrated stress response (ISR) in SCC cells, we generated eIF2α-S51A, or “ISR-null” cancer cells using a knockout and reconstitution strategy. For this purpose, we used an aggressive HRas^G12V^ murine, primary-derived skin SCC line expressing an eGFP reporter (Yang et al., 2015). We transduced these cells with lentiviral constructs harboring a PGK promoter-driven cDNA encoding either the wild-type (S51) or phospho-dead (S51A) eIF2α protein (encoded by the *eif2s1* locus) (Figure 1B). The *eif2s1* transgenes each contained a synonymous mutation in a protospacer adjacent motif (PAM) site that rendered it resistant to a small guide RNA (sgRNA) that could be used to specifically target the endogenous *eif2s1* gene for CRISPR/Cas9 deletion.

After cells were transduced with the myc-tagged *eif2s1* constructs, cells were transfected with liposomes harboring CRIPSR-Cas9/sgRNA ribonucleoproteins, which targeted the ablation of the endogenous *eif2s1* alleles, and thus left the myc-tagged S51 or S51A transgenes as the sole source of eIF2α expression. Following ablation and reconstitution, single cells were isolated by fluorescence activated cell sorting (FACS) and used to generate stable SCC clones. Successful targeting of the two endogenous *eif2s1* alleles was verified by genomic sequencing (Figure 1B).

Two of each S51 and S51A eIF2α replacement clones were chosen for further study. In all assays presented in Figures 1 and 2, the clones of the same eIF2α status grew and behaved similarly. Hence for the purposes of the current study, we show the results on pooled clones displaying common genotypes. Immunoblot analyses revealed that total eIF2α levels were comparable to the parental eIF2α clone, and the replacement eIF2α proteins exhibited the expected increase in size due to the epitope tag (Figure 1C). Importantly, when we treated these SCC lines with sodium arsenite to induce oxidative stress and activate the HRI kinase (Sendoel et al., 2017), we observed that like the parental clone, the stressed eIF2α S51 cells displayed phospho-S51 immunolabeling, while the eIF2α S51A clones were refractory to phosphorylation. Taken together, these results underscored the efficacy of our knockout and replacement strategy, and verified the dramatic difference in stress-induced abilities of our two clones to target eIF2α phosphorylation at the heart of the ISR.

**Figure 2:**
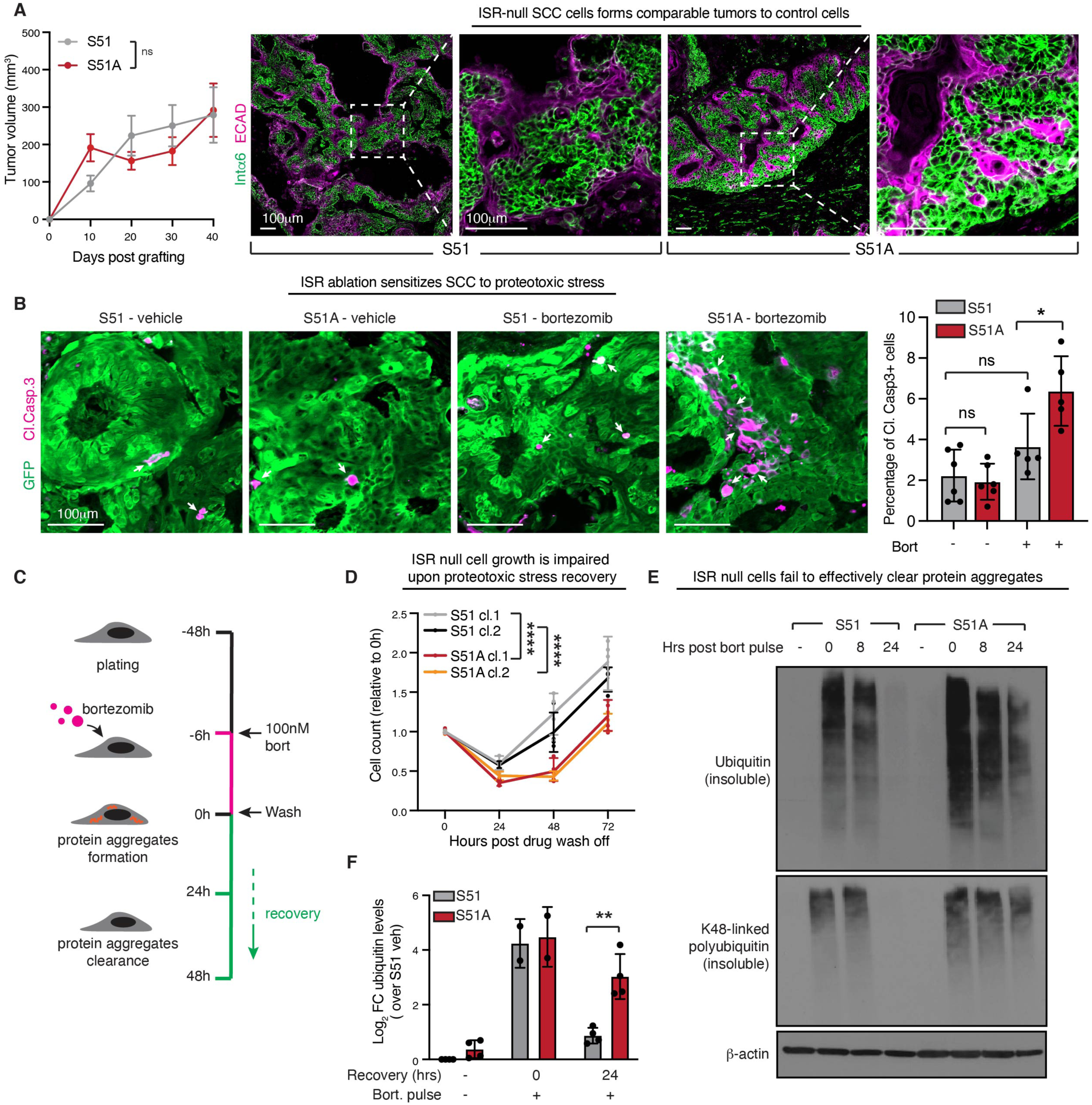
The ISR is required for efficient clearance of protein aggregates. A. Left, growth of tumor allografts derived from control or ISR-null SCC cells. Tumor volume is comparable between conditions. Error bars represent SEM from N=16 tumors per condition. ns, no statistical significance (simple linear regression). Right, α6-integrin and E-cadherin immunofluorescence of SCC at day 17 reveals that ISR-ablation does not impact tumor morphology. Boxed regions are shown at higher magnification images to the right of each image. Scale bar 100 μm. B. Cleaved caspase 3 immunofluorescence of tumor sections reveals elevated apoptosis, a sign of increased sensitivity of ISR-null SCC tumors to proteotoxic stress. (Bortezomib treatment 1.2 mg/kg i.p. on days 13, 15, and 17). No significant difference in viability is observed in vehicle treated SCC tumors. White arrows indicate cleaved caspase-3 positive cells. Bar graph on the right shows percentage of cleaved caspase-3 positive cells ± SD within the GFP+ tumor cells, in at least 5 independent tumors. * P<0.05, ns, no statistical significance (one-way ANOVA, multiple comparisons). Scale bar 100 μm. C. Schematic of protein aggregate pulse-recovery experiments. Cells are treated with a saturating (100 nM) dose of bortezomib for 6 hrs, a timepoint at which protein aggregates have accumulated, but cells are still viable. Bortezomib is washed from the media and cells are then monitored during the recovery phase. D. Cellular proliferation during the protein aggregate recovery phase indicates that control cells regain proliferative capacity 24 hrs prior to ISR-null cells. N=4 biological replicates, n=16 total technical replicates, error bars denote SD of technical replicates. ****P<0.0001 (two-way ANOVA, multiple comparisons). E. Protein aggregates evaluated by anti-ubiquitin and K48-linked polyubiquitin immonblot following cellular fractionation of soluble and insoluble proteins. Representative immunoblot of the insoluble fraction shows clearance of aggregates in control but not ISR-null cells at 24 hrs. F. Bar graph shows log2 fold change (FC) ubiquitin levels ± SD normalized over *β*-actin and relative to vehicle-treated control cells. **P<0.01 (t test).

To further interrogate the functionality of our SCC lines in mounting an ISR, we monitored their protein synthesis rates in response to stress. Using puromycin incorporation as a gauge to measure nascent protein synthesis, we observed that translation within eIF2α S51 cells was comparable to the native eIF2α parental clone, and markedly dampened upon stress as expected (Figure 1D). Consistent with the inability of eIF2α-S51A to become phosphorylated and decommissioned, the eIF2α-S51A SCC cells sustained higher levels of protein synthesis during oxidative stress than their control counterparts (Figure 1D). Similar results were seen upon exposure of our SCC cell lines to tunicamycin, which induces ER-stress and eIF2α-S51 phosphorylation through PERK kinase.

Proteosome inhibitors such as the boronic acid dipeptide derivative, bortezomib, are known to result in accumulation of misfolded proteins, ER stress and eIF2α phosphorylation (Jiang & Wek, 2005; Suraweera et al., 2012; Yerlikaya & Okur, 2020). Typical of a classical ISR, global protein synthesis was reduced in our SCC cells exposed to bortezomib, consistent with pathway activation (Figure 1D and Figure 1-figure supplement 1A). In contrast, however, there did not seem to be a marked difference in protein synthesis rates between bortezomib treated eIF2α-S51 and eIF2α-S51A cells. In this regard, SCC cells appeared to face proteotoxic stress in a manner distinct from a classical integrated stress response.

Probing deeper, we found that under the serum-rich culture conditions used here, S51 and S51A cells proliferated at comparable rates and showed no differences in viability (Figure 1-figure supplement 1B-C). Analogously, the morphologies of these cells were indistinguishable under these conditions (Figure 1-figure supplement 1D). However, upon exposure to stress, eIF2α-S51A cells were significantly more sensitive than SCC cells with an intact ISR. Particularly, after 24 hrs of bortezomib treatment, the percentage of cells positive for the apoptotic marker Annexin-V was appreciably higher in S51A over S51 cultures (Figure 1E; Figure 1-figure supplement 1E). Taken together, our collective data thus far pointed to the view that when faced with proteotoxic stress, SCC cells harboring a functional ISR may gain a fitness advantage over their ISR null counterparts not by selectively dampening down global protein synthesis, but rather by shifting to a translational landscape that is more favorable for survival.

### The ISR promotes efficient clearance of protein aggregates

When epidermal stem cells acquire a single oncogenic mutation and will eventually progress to SCC, they activate alternative translational pathways suggestive of an ISR (Blanco et al., 2016; Sendoel et al., 2017). Intriguingly, however, when we injected our SCC cells into the skins of immunocompromised, athymic (*Nude*) mice, S51A and S51 SCC cells both formed tumors that grew similarly and showed similar morphologies (Figure 2A; Figure 2-figure supplement 1A). This was not attributable to ‘escapers’ that had somehow circumvented the S51A mutation *in vivo*, as FACS sorted SCC stem cells from our S51A tumors still displayed insensitivity to sodium arsenite-induced eIF2α phosphorylation in contrast to their S51 counterparts (Figure 2-figure supplement 1B).

Most strikingly, when we treated our mice with bortezomib, the *in vivo* S51A SCC tumors exhibited significantly greater sensitivity to proteotoxic stress and apoptosis than their counterparts (Figure 2B). These findings were particularly relevant given that despite intense interest in proteasome inhibitors as a potential new line of cancer therapeutics, solid tumors have shown resistance to these drugs (Fournier et al., 2010; Manasanch & Orlowski, 2017). Taken together, our results raised the tantalizing possibility that if the ISR is first crippled in solid tumors, their tumor-propagating stem cells may be increasingly sensitive to added stress.

While the ISR is known to be activated by the types of unfolded proteins that accumulate upon proteasome inhibition (Figure 1-figure supplement 1A)(Pakos-Zebrucka et al., 2016), it is incompletely understood how the pathway promotes cell recovery and survival in the face of this stress. Intrigued that ISR-null cells and tumors had an intrinsic liability in coping with protein aggregate stress, we established a two-step *in vitro* model of first triggering SCC cells to accumulate an excess of unfolded proteins and then allowing them to recover from the stress. To this end, we treated cultured control or ISR-null SCC cells *in vitro* with a saturating dose of bortezomib for 6 hrs, at which point we washed the cells and switched to fresh media to allow cells to recover (Figure 2C). Following this bortezomib “pulse”, control SCC cells with a competent ISR recovered and began to proliferate within 24 to 48 hrs. In striking contrast, ISR-null SCC cells took a full 24 hrs longer than their counterparts before they began to proliferate again (Figure 2D). Intriguingly, this could not be imputed to differences in viability, since following this treatment regime, ISR-null and ISR-intact SCC stem cells displayed no difference in Annexin-V positivity either at 6 hrs of bortezomib treatment or at 24 hrs after recovery (Figure 2-figure supplement 2A-C).

The first step in targeting unfolded proteins for degradation is their ubiquitination (Smith et al., 2011). We therefore examined the clearance of ubiquitinated proteins in SCC cells recovering from proteotoxic stress. Following treatment with bortezomib, cells were lysed in Radio-Immunoprecipitation Assay (RIPA) buffer, and soluble and insoluble proteins were then fractionated by centrifugation. Each fraction was normalized based on the protein concentration of the soluble fraction and then subjected to polyacrylamide gel electrophoresis and analyzed by anti-ubiquitin immunoblots.

ISR-null and control SCC cells responded similarly to bortezomib, displaying a rapid jump in ubiquitinated proteins within 6 hrs of treatment (Figure 2E-F). After withdrawing bortezomib, however, the rate at which ubiquitinated proteins were cleared from the insoluble fraction was remarkably reduced in the ISR-null cells compared to controls SCC cell lysates. As lysine 48-linked polyubiquitin is the specific mark of proteins targeted for proteasomal degradation (Thrower et al., 2000), we performed anti-K48-polyubiquitin immunoblot analyses (Figure 2E). These data confirmed that SCC cells that are unable to mount an ISR are not defective in their E3-ubiquitin-ligase system per se, but rather are impaired in their ability to efficiently clear proteins that are marked for proteosomal destruction.

We also used this assay to interrogate the source of the insoluble protein aggregates induced by bortezomib. To do so, we treated cells concurrently with cycloheximide, to block new protein synthesis and with bortezomib, to inhibit proteosome-mediated aggregated protein clearance. Inhibiting translation elongation with cyclohexamide or translation initiation with harringtonine nearly quantitatively blocked the buildup of protein aggregates following 6 hrs of bortezomib treatment. These data suggested that if an acute block in proteasome function occurs and SCC cells cannot respond quickly by slowing new protein synthesis, an imbalance in proteostasis arises, resulting in an accumulation of newly synthesized, unfolded proteins (Figure 2-figure supplement 3A).

We next addressed the extent to which the proteosome versus autophagy is responsible for the clearance of these ubiquitin-marked protein aggregates during the recovery phase following bortezomib treatment. To do so, we treated with bortezomib for 6 hrs, and then after washing out the drug, we either allowed both pathways to participate in clearance or added bafilomycin A1 to block the autophagy pathway. As shown in Figure 2-figure supplement 3B, BafA1 delayed recovery partially but not fully. Taken together, these data indicated that both autophagy and the proteasome cooperate in clearing these ubiquitin-marked protein aggregates during the recovery phase following proteotoxic stress.

### The ISR responds to proteotoxic stress by upregulating translation of centrosomal proteins

Our findings pointed to the view that the ISR is a major player in the process of cellular recovery from proteotoxic stress. The delayed rate in protein aggregate clearance seen in ISR-null cells was not attributable to a higher rate of global protein synthesis, as this was comparable to the ISR-competent control cells (Figure 1D). We therefore asked whether the ISR might be required to drive translation of select mRNAs upon proteotoxic stress, and if so, whether the proteins produced under such circumstances might give us clues into how the ISR functions in recovery. To this end, we performed ribosome profiling to landscape the ISR-mediated impact on translation during proteotoxic stress (Figure 3A) (Ingolia et al., 2009; McGlincy & Ingolia, 2017).

**Figure 3.**
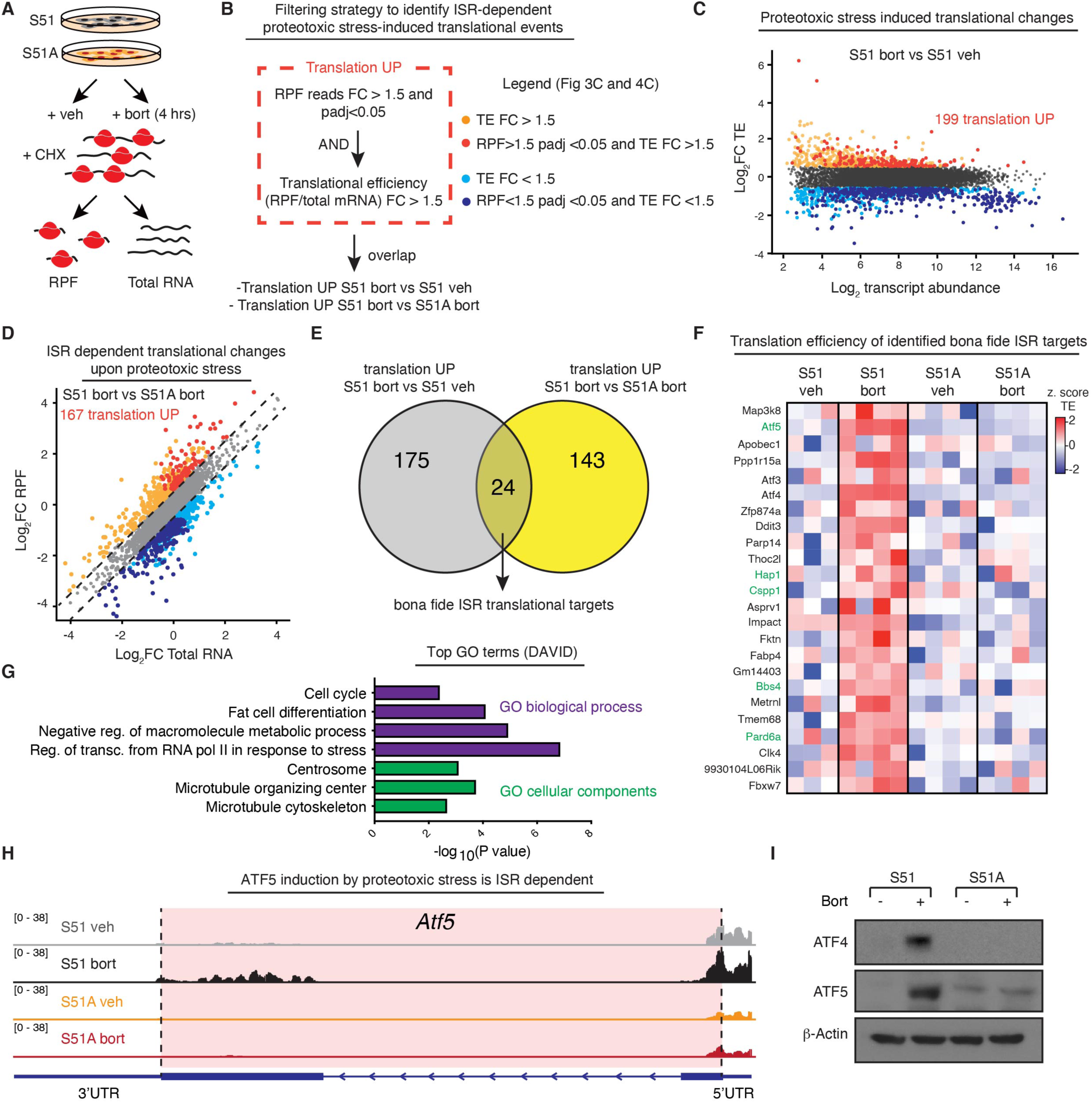
The ISR induces synthesis of centrosomal proteins in response to proteotoxic stress. A. Schematic of ribosome profiling experiment. N=4 samples per condition of control or ISR-null cells were treated with or without 100 nM bortezomib for 4 hrs. Cells were lysed in the presence of 100 μg/mL cyclohexmide and total mRNA and ribosome protected fragments were sequenced to analyze translational efficiency genome-wide. B. Filtering strategy to identify a high-confidence list of translationally-regulated mRNAs targeted by the ISR following protein aggregate stress. To identify translationally regulated mRNAs, a two-step approach was taken. First, genes with ribosome protected fragments (RPF) of a fold change > 1.5 and padj <0.05 (DESeq2 generalized linear model) were identified. Second, within this list, genes with translational efficiency (TE) fold change > 1.5 were selected (red dots, panel C and D). Finally, ISR target genes are identified by the overlap of mRNAs with significantly upregulated translation in two comparisons: (1) ISR-competent-bort vs. ISR-competent-vehicle and (2) ISR-competent-bort vs. ISR-null-bort. C. Dot plot shows translational efficiency log_2_ fold change vs. log_2_ mean transcript abundance genome-wide in ISR-competent-bort vs. ISR-competent-vehicle highlights both up-regulated genes as well as a down-regulated group of highly abundant transcripts (see legend in panel B). As expected, highly expressed house-keeping transcripts are translationally downregulated. D. Dot plot shows RPF log_2_ fold change vs. total mRNA log_2_ fold change genome wide in ISR-competent-bort vs. ISR-null-bort (see legend in panel B). E. Pie chart shows high confidence translational up-regulated ISR targets. Overlap between translationally upregulated genes in (1) ISR-competent-bort vs. ISR-competent-vehicle and (2) ISR-competent-bort vs. ISR-null-bort is shown. F. Z-scored heat map of translational efficiencies for 24 genes translationally induced by the ISR. Centrosomal proteins are highlighted in green lettering. G. Top gene ontology (GO) terms identified in categories of biological processes and cellular components using translationally induced genes by the ISR. Centrosomal proteins are clearly enriched among cellular components. H. Representative RPF tracks for *Atf5* reveal coding sequence translation is specifically induced in control cells treated with bortezomib. *Atf5* coding sequence is shaded in red. I. Representative immunoblot confirms ISR-dependent induction of ATF4 and ATF5 following protein aggregate stress.

Briefly, we subjected control and ISR-null SCC cells to proteotoxic stress (bortezomib) versus vehicle control. Quadruplicate samples of each condition were then lysed in the presence of cycloheximide in order to preserve the ribosome location along transcripts. Total mRNAs were saved, and the remaining lysates were treated with RNase-I to digest away all mRNAs that were not protected by ribosomes. Total mRNAs and the ribosome protected fragments (RPFs) were then prepared for deep sequencing, and the translational efficiency was analyzed by assessing the ratio of RPFs to total mRNA reads, genome wide.

Reads were aligned to the mouse reference genome, and quality control was performed to confirm that we had successfully purified and sequenced RNA fragments that were protected by actively translating ribosomes. Technical replicates were found to co-vary within samples by subjecting them to principal component analysis (Figure 3-figure supplement 1A). As a second quality control, metagene analysis was performed, which demonstrated that the majority of RPFs were found within coding segments, with relative peaks at the translation start and stop sites, as would be expected from a high quality ribosome profiling dataset (Figure 3-figure supplement 1B). The read lengths peaked between 29 and 32 nucleotides, corresponding to the correct length of mRNA protected by ribosomes from RNase-I digestion (McGlincy & Ingolia, 2017) (Figure 3-figure supplement 1C).

Having verified the efficacy of our data, we next focused on the translational response to proteotoxic stress in eIF2α-S51 control SCC cells bearing an intact ISR. To this end, we sought to identify mRNAs that displayed increased ribosome occupancy (RPF FC >1.5 and padj(RPF) <0.05) in bortezomib compared to vehicle treated cells. To eliminate possible translational variances arising from transcriptional differences, we also normalized the RPF reads of each transcript according to total mRNA levels (Ingolia et al., 2009; McGlincy & Ingolia, 2017). This allowed us to identify genes with translational efficiencies (TE) (RPF/mRNA) that are sensitive to bortezomib, and whose translational changes are at the heart of ISR-mediated differences (Figure 3B). Interestingly, in our S51-SCC cells with an intact ISR, proteotoxic stress provoked the translational upregulation of 199 mRNAs (RPF FC >1.5, padj (RPF) < 0.05 and TE FC > 1.5) (Figure 3C).

Next, we performed the same analysis to compare ISR-competent and ISR-null cells upon proteotoxic stress. This revealed 167 mRNAs whose translational upregulation was dependent on the presence of an intact ISR (S51-bort vs S51A-bort (RPF FC >1.5, padj (RPF) < 0.05 and TE FC > 1.5)) (Figure 3D). These findings clearly showed that when SCC cells are refractory to proteotoxic stress-induced phosphorylation of eIF2α, their translational program is selectively perturbed.

To curate a list of specific ISR-targeted mRNAs, we identified genes for which the translation changed in response to bortezomib and only in cells with an intact ISR. To this end, we generated a Venn Diagram, comparing (A) the 199 mRNAs translationally upregulated in control cells following bortezomib treatment, and (B) the 167 mRNAs that failed to be translationally upregulated by bortezomib when the ISR was crippled. By this criterion, 24 mRNAs surfaced as candidates for *bona fide* ISR translational targets and showed strong enrichment across all four sets of translational replicates of proteotoxic stress-induced SCC cells harboring an intact ISR (Figures 3E and 3F).

We used gene ontology enrichment analysis (GO-term) to ask if the ISR targeted sets of mRNAs corresponded to common biological processes or cellular components. The downregulated genes were clearly enriched in components of translational machinery, which is consistent with the known role for the ISR pathway in downregulating translation and housekeeping gene synthesis (Figure 3-figure supplement 1D). GO-term analysis of the 24 upregulated ISR-targets was especially interesting (Figure 3G). Regulation of stress induced transcription featured prominently in the top GO-terms for biological processes and was exemplified by the presence of ATF4, a well-established stress induced transcriptional regulator that is activated in premalignant SCC cells by non-canonical translation when eIF2α is phosphorylated (Sendoel et al., 2017). Most intriguing, however, were the top three GO-terms for cellular components: Centrosomal proteins, Microtubule organizing center (MTOC) proteins and Microtubule cytoskeletal proteins (Figures 3G and genes highlighted in green in Figure 3F).

The centrosome is the major cytoplasmic MTOC within interphase cells (Caviston & Holzbaur, 2006; Sanchez & Feldman, 2017; Woodruff et al., 2017). In SCC cells, like most other mammalian cells, the MTOC is located near the nucleus, where it is surrounded by pericentriolar proteins important for plus-ended microtubule growth towards the cell periphery. The polarity of microtubules establishes polarized transport, which depending upon the motor protein involved, can occur either towards the minus ends at the MTOC, or along the plus ends of the microtubules towards the focal adhesions.

Although the ISR had not been previously found to regulate the microtubule cytoskeleton, we were intrigued by the possibility that the ISR was influencing microtubule dynamics in response to proteotoxic stress, which in turn might affect how efficiently proteins are cleared during the recovery phase. This possibility was all the more compelling because protein aggregate formation and clearance is known to depend on microtubule-dependent intracellular transport (Kopito, 2000). Of further intrigue was the mRNA for ATF5, which like that for ATF4, displayed dramatic increases in translation specifically in ISR-competent SCC cells exposed to proteotoxic stress and not in the ISR-null SCC counterparts (Figures 3H and 3I). ATF5 has been previously demonstrated to play a non-transcriptional role at the centrosome(Madarampalli et al., 2015), strengthening the hypothesis that ISR activation was driving a subset of centrosomal proteins to preserve the response to proteotoxic stress. We therefore set out to evaluate how the centrosome and microtubule dynamics were changing in response to stress in SCC cells with or without an intact ISR.

### The ISR protects centrosomal microtubule dynamics

To evaluate the status of the centrosome during stress, we performed immunofluorescence for centrosomal markers pericentrin (PCNT) and γ-tubulin (TUBG1). Fluorescence intensities of both markers were significantly increased in control, but not ISR-null, cells and specifically in response to proteotoxic stress (Figure 4A and Figure 4-figure supplement 1). Quantifications further revealed that the overall size of the MTOC was also enlarged in response to this stress, but only when the ISR was intact (Figure 4B). This was especially intriguing because the pericentriolar region that surrounds the centrioles of the microtubule organizing centers is a special site of protein catabolism within the cell, concentrating both autophagosome and lysosomes as well as proteasomes and proteins marked for degradation (Freed et al., 1999; Liu et al., 2016; Wigley et al., 1999) . Moreover, although the highly transformed, long-passaged HeLa cell line lacks physiological relevance, it was notable that proteosome inhibition in these cells resulted in the accumulation of pericentriolar material, including pericentrin and γ-tubulin (Didier et al., 2008). When we placed these data in the context of our new findings, we became all the more interested that the ISR may be at the crux of reinforcing microtubule dynamics following proteotoxic stress.

**Figure 4:**
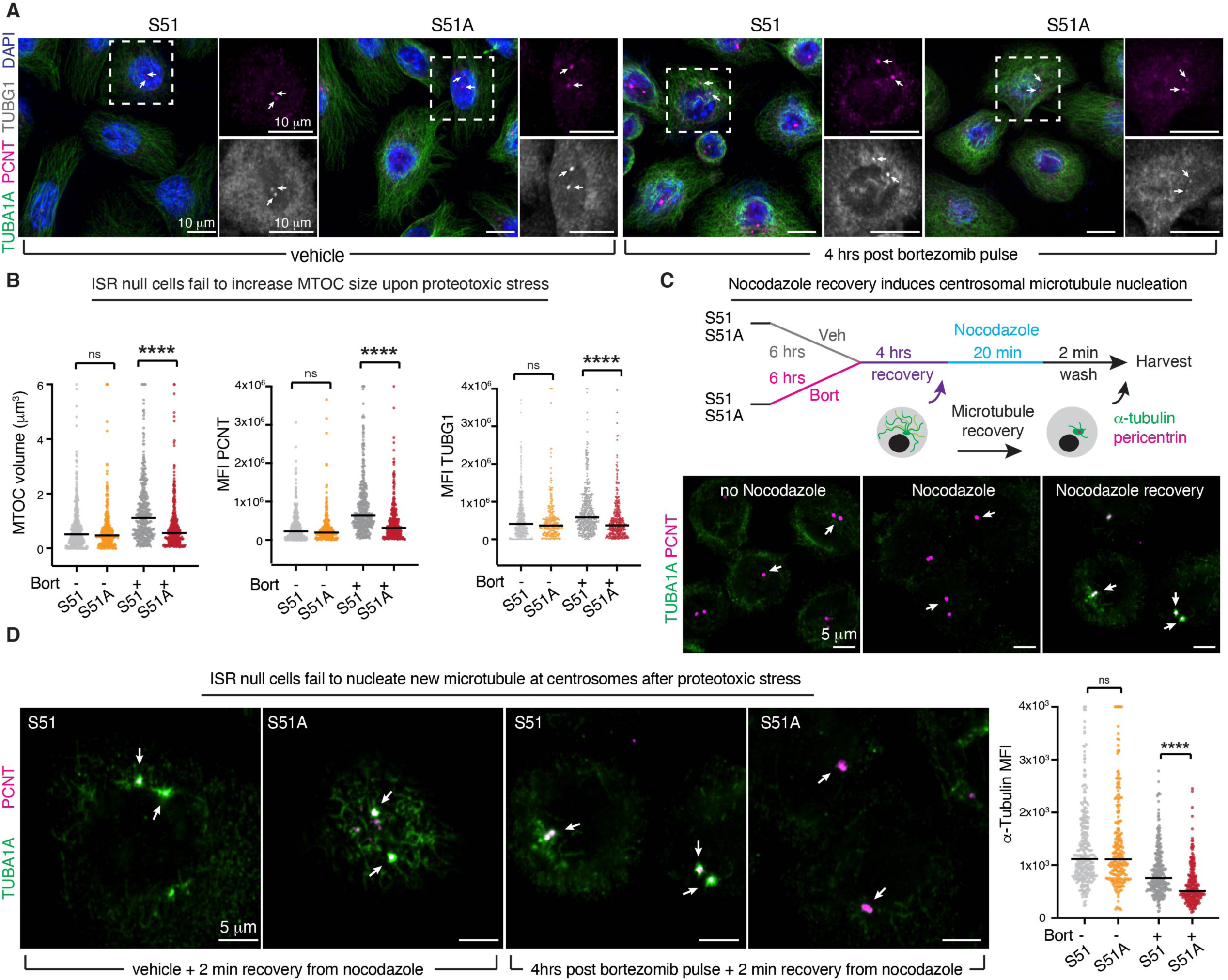
The ISR protects centrosomal microtubule dynamics during recovery from protein aggregate stress. A and B. Representative immunofluorescence images of S51 and S51A cells vehicle-treated or treated with a bortezomib pulse (100 nM, 6 hrs) and let to recover for 4 hrs. Immunolabeling is to highlight the centrosomal MTOC (pericentrin, PCNT and γ-tubulin, TUBG1), and the cytoplasmic microtubules (α-tubulin, TUBA1A). Scale bar 10 μm. White arrows indicate centrosomal MTOCs. B. Both volume and fluorescence intensity of MTOC markers are increased specifically in control cells recovering from protein aggregate stress. All data are visualized on a per centrosome basis from N=2 independent experiments. ****p<0.0001 ns, no statistical significance (two-way ANOVA with multiple comparisons). C. Schematic of microtubule recovery assay to assess the capacity of centrosomes to nucleate new microtubules following 4 hrs recovery from protein aggregate stress. Cells with or without protein aggregates are treated with 13 μM nocodazole for 20 min to depolymerize microtubules. Nocodazole is washed from the media, and new microtubules are allowed to form during a 2 min incubation at room temperature. Cells are fixed and MTOCs (PCNT+) and cytoplasmic microtubules (TUBA1A) are quantified and analyzed by immunofluorescence. Bottom, representative images demonstrate complete depolymerization of microtubules by nocodazole and nucleation of new centrosomal microtubules following nocodazole wash. White arrows indicate centrosomal MTOCs. Scale bar 5 μm. D. Representative images of centrosomal microtubule nucleation following nocodazole wash out in control and ISR-null cells with or without protein aggregates. Centrosomal microtubule nucleation is analyzed by measuring α-tubulin signal at the centrosome, which is defined as the pericentrin-positive volume. White arrows indicate centrosomal MTOCs. Scale bar 5 μm. Quantification of mean centrosomal α-tubulin intensity indicates that control cells are partially protected against impaired microtubule dynamics during stress. N=2 independent experiments. ****p<0.0001 ns, no statistical significance (two-way ANOVA with multiple comparisons).

If our premise was valid, then when cells cannot trigger an ISR in the face of stressful situations, microtubule-dependent processes should be vulnerable. To evaluate microtubule dynamics during the recovery phase of our cells following proteotoxic stress exposure, we briefly interrupted existing microtubule assembly/disassembly dynamics with nocodazole and then examined nascent microtubule nucleation initiated from the centrosome (Figures 4C and 4D).

In the absence of stress, the ISR-null state did not appreciably affect MTOC-initiated microtubule dynamics (Figure 4D). In striking contrast, the vulnerability of microtubule dynamics in the ISR-null state became clear when we exposed our cells to the proteotoxic stress and monitored the recovery process. Even after 4 hrs of bortezomib recovery, microtubule growth from the MTOC was still markedly impaired in the ISR-null compared to control SCC cells (Figure 4D). Since microtubule dynamics were independent of the ISR in the unstressed state, these results suggested that the ISR is required to reinforce the microtubule dynamics involved in cellular recovery following proteotoxic stress.

### The ISR is required for aggresome formation

Our evidence that protecting microtubule dynamics during proteotoxic stress is a major function of the ISR was all the more compelling because of known role for microtubules in assembling ubiquitinated protein aggregates into larger membraneless structures called aggresomes (Johnston et al., 1998; Wigley et al., 1999). Through their ability to associate with proteasomes and autophagosomes, aggresomes appear to be a key intermediate in the clearance of improperly folded proteins.

With these insights in mind, and the general view that aggresomes function beneficially in cellular recovery from proteotoxic stress, we evaluated aggresome formation in our SCC cells following withdrawal of bortezomib (Figure 5A). To this end, we used immunofluorescence for p62/SQSTM1, a protein involved in shuttling ubiquitinated protein aggregates to the aggresome (Christian et al., 2010). As judged by convergence of intense anti-p62 immunofluorescence to a single large perinuclear spot within each S51 cell, misfolded protein aggregates began to coalesce into the aggresome soon after bortezomib withdrawal, peaking at approximately 8 hrs into the recovery phase (Figure 5B, top panels). By 24 hrs, the recovery phase appeared to be complete, as the aggresome was no longer present (see quantifications at right). These findings were in good agreement with the clearance of ubiquitinated misfolded proteins that had accumulated during the proteosome block (Figure 2E).

**Figure 5:**
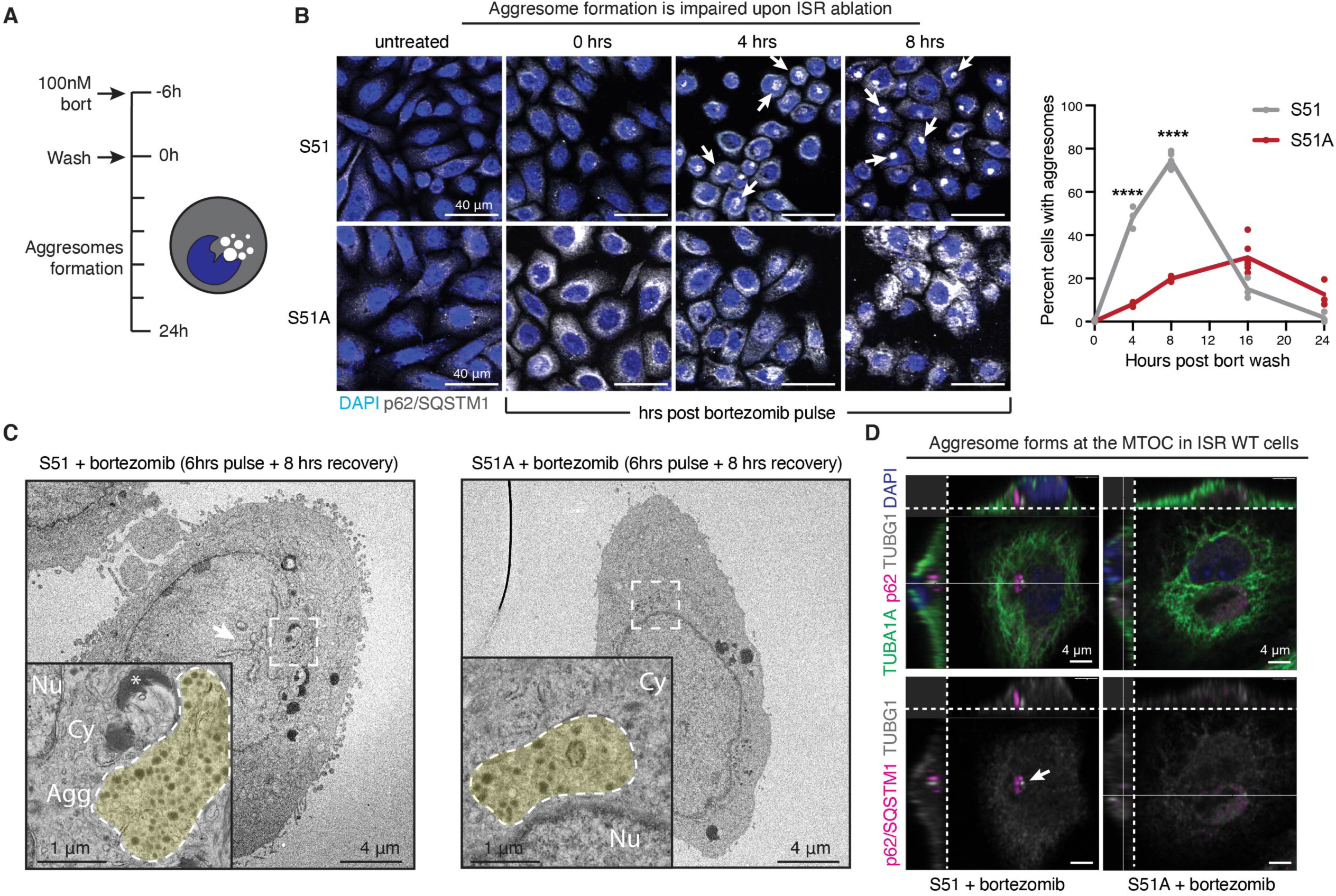
The ISR is required for aggresome formation. A. Schematic of aggresome formation assay. During recovery from proteotoxic stress, aggresome formation is assessed by anti-p62/SQSTM1 immunofluorescence, which appears as a well-defined, perinuclear puncta. B. Time course of aggresome formation in control and ISR-null cells indicates diminished aggresome formation in cells with an ablated ISR. White arrows indicate aggresomes. Quantification of the percentage of aggresome-positive cells. N=2 independent experiments with at least 145 cells quantified per condition. Error bars represent 95% confidence interval calculated by Wilson’s method for confidence interval of a proportion. ****p<0.001 using two sample Z-test for a proportion (comparisons: control vs. ISR-null at 4 hrs and 8 hrs). Scale bar 40 μm. C. Aggresomes visualized at the ultrastructural level by transmission electron microscopy. In control cells aggresomes characteristically appear as a perinuclear cluster of electron dense granules and partially-degraded organelles that often deform the nuclear membrane. Fewer aggresomes form in ISR-null cells, and those that do are smaller, more diffuse and rarely indent the nucleus. White arrow indicates nuclear envelope deformation. Scale bar 4 μm. Yellow shade in the zoomed-in images at the lower left highlights aggresomes and protein aggregates. Nu, nucleus. Cy, cytoplasm, Agg, aggresome. *, lipid droplet. Scale bar 1 μm. D. Confocal immunofluorescence microscopy and 3D reconstruction to visualize the location of the aggresome in relation to the MTOC (γ-tubulin, TUBG1) and cytoplasmic microtubules (α-tubulin, TUBA1A). In control cells, aggresomes appear as dense p62-positive structures directly adjacent to the MTOC puncta. Representative cells from 8 hrs post-bortezomib wash out are shown. White arrow indicates MTOC. Scale bar 4 μm.

In striking contrast to S51 cells, the S51A cells lacking an intact ISR showed pronounced defects in their ability to clear misfolded proteins that had accumulated during bortezomib treatment. The rise in cytoplasmic p62 immunofluorescence during the proteosomal block indicated that small protein aggregates had formed in the S51A cells exposed to proteotoxic stress (Figure 5B, bottom panels). However, even at 8 hrs after bortezomib withdrawal, only a few S51A cells displayed the bright perinuclear spot reflective of the aggresome. Moreover, while S51 cells had cleared their protein by 24 hrs after culturing in normal media, S51A cells still exhibited anti-p62 coalescence, indicative of a marked delay in clearance of unfolded proteins.

Further signs of cellular defects during the recovery process were evident at the ultrastructural level. After 8 hrs of recovery, electron dense protein aggregates pushing into the nucleus were observed in the perinuclear regions of S51 but not S51A cells. These structures were absent in unstressed cells. (Figures 5C and Figure 5-figure supplement 1A-C). Co-immunolabeling for p62 and γ-tubulin confirmed that the perinuclear structures present in S51 cells were indeed aggresomes (Figure 5D). Moreover, S51 cells displayed signs of nuclear deformation, often observed during the aggresome clearance phase. By contrast, when detected in S51A cells, aggresomes tended to be small. Rather, S51A cells often displayed widespread vacuole-like structures which have been reported to occur when the ER becomes overwhelmed with misfolded proteins (Mimnaugh et al., 2006) (Figure 5-figure supplement 1C).

Since these structures did not accumulate in S51 cells, it seemed likely that the root of this phenotype was the inability of ISR-null cells to cope with *de novo* protein synthesis in the face of acute proteotoxic stress. However, this did not explain why clearance of cytoplasmic protein aggregates was markedly diminished in S51A cells. Rather, the surprising dependence upon an intact ISR for SCC cells to form aggresomes during proteotoxic stress recovery led us to posit that the ISR functions critically in sustaining the necessary microtubule dynamics to enable efficient transport of misfolded proteins to the MTOC, where they can form aggresomes and be targeted for perinuclear proteosomal and autophagosomal clearance.

To further challenge the relation between microtubule dynamics and aggresome formation, we used the drug, paclitaxel, a chemotherapy that stabilizes microtubules while at the same time disrupting their dynamics. Strengthening the link between the ISR, microtubule dynamics, and aggregate clearance, treating cells with paclitaxel following the induction of protein aggregates potently blocked aggresome formation, phenocopying the ISR-null cells during this process (Figure 5-figure supplement 2A). Blocking microtubules polymerization by treating cells with nocodazole had a similar effect and resulted in a significant reduction in aggresome formation (Figure 5-figure supplement 2B).

### The integrated stress response is required for migration and focal adhesion homeostasis following protein aggregate stress

During the course of our previous experiments, we made several observations regarding the cellular morphology, which supported the conclusion that the ISR was required to maintain proper microtubule dynamics in the face of protein aggregate stress. First, we noticed that control, but not ISR-null cells, went through a dramatic change in cell shape while recovering from aggregate stress. Specifically, control cells “unspread”, becoming more compact and tall between 4 and 8 hrs after washing out bortezomib, the timepoints correlating with peak aggresome formation. ISR-null cells on the other hand, did not similarly round, and instead they maintained a flattened shape with some cells also forming elongated processes. Examples of these differences are shown in Figure 6-figure supplement 1.

The long cellular extensions were reminiscent of that seen when keratinocytes displayed defects in the turnover of focal adhesions, a process that we had previously shown depends not only upon focal adhesion proteins (Schober et al., 2007), but also on the ability of microtubules to deliver turnover cargo to the focal adhesions (Wu et al., 2008). Specifically, microtubule-mediated cargo transport is essential for focal adhesion disassembly and cell migration (Ezratty et al., 2005; Yue et al., 2014). While the transport of protein aggregates to the centrosome involves minus end-directed molecular motors, transport to the focal adhesions requires plus end-directed motors. Reasoning that a defect in MTOC-mediated microtubule dynamics might affect both processes, we examined the focal adhesions in control and ISR-null cells following proteotoxic stress. As judged by immunofluorescence for vinculin, upon proteotoxic stress the ISR promoted a remodeling in cell morphology evident by diminished focal adhesion and F-actin staining. By contrast, ISR-null cells retained large focal adhesions suggestive of impaired cytoskeletal dynamics. (Figure 6A). This ISR-mediated difference was specific to stress and was not seen when control and ISR-null cells were cultured in normal media under unstressed conditions. Compellingly, further analysis of our ribosome profiling dataset revealed a significant enrichment of focal adhesion genes within mRNAs translationally downregulated upon ISR induction (Figure 3-figure supplement 1D), suggesting that ISR-mediated translational control stands at the crux of these dramatic morphological changes by selectively upregulating microtubule dynamics and, at the same time, downregulating focal adhesion components.

**Figure 6:**
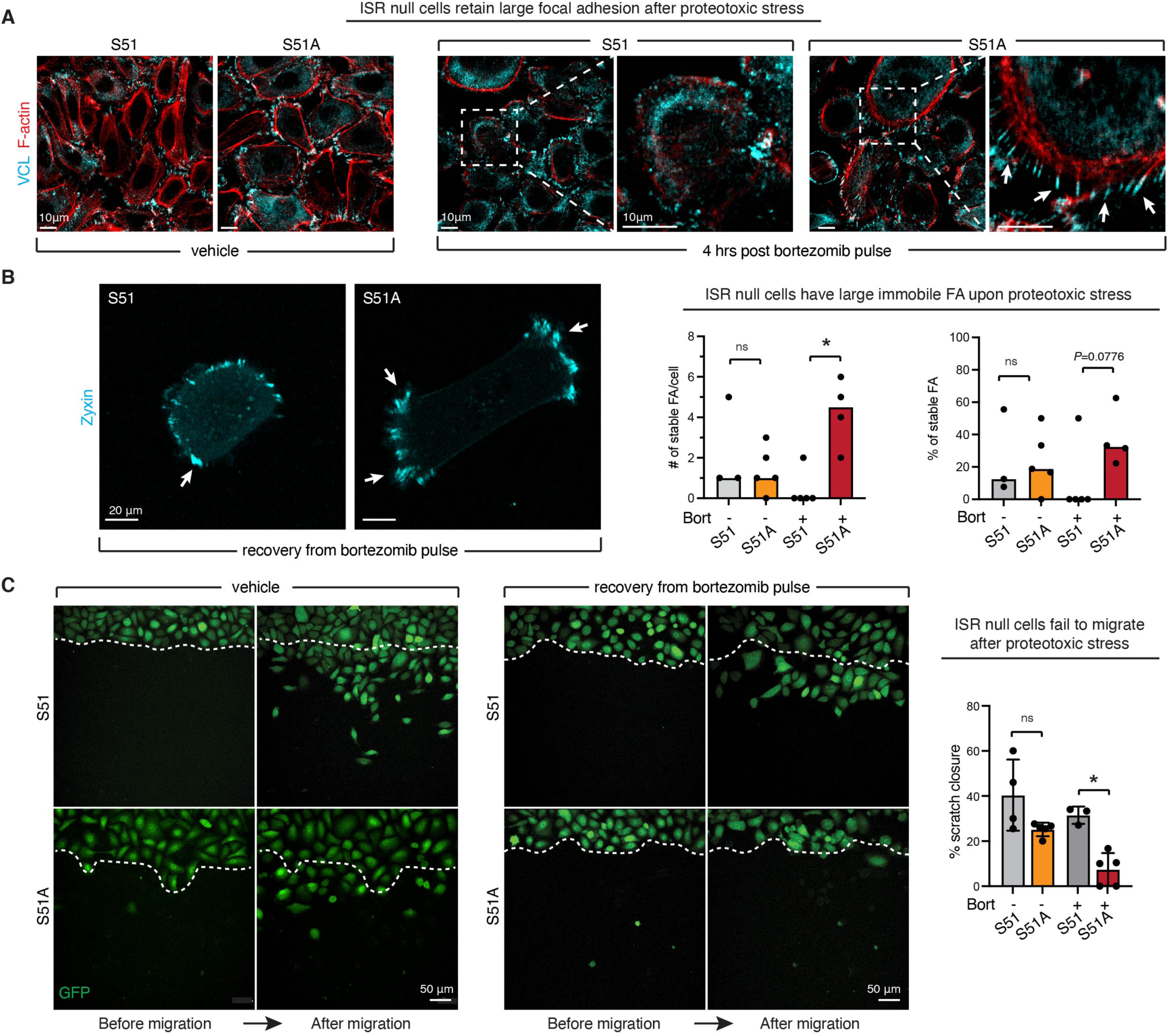
The ISR is required for focal adhesion remodeling and cell migration upon proteotoxic stress. A. Impaired focal adhesion dynamics in ISR-null cells. Immunofluorescence for vinculin and F-actin shows that during recovery from protein aggregate stress, the ISR promotes the remodeling of focal adhesions. White arrows show characteristic long cellular extensions connected to enlarged focal adhesions in the ISR-null but not control cells recovering from protein aggregate stress. Consistent with the known role of microtubules in targeting focal adhesion turnover, these features resemble those of keratinocytes lacking microtubule-focal adhesion targeting proteins (Wu et al., 2008). Scale bar 10 μm. B. Static frames from live imaging of zyxin-iRFP cells. White arrows show large (10 μm^3^ or greater) and stable (over the whole duration of imaging period) focal adhesion clusters. Bar graphs show total number of stable and large focal adhesion clusters per cell (left) and percentage of stable over total focal adhesion clusters (right) over a 14 min live imaging time course. 3-5 cells were imaged per condition in two independent biological replicates. *p<0.05 ns, no statistical significance (t test). C. Live imaging of GFP-positive cells migrating in a scratch assay reveals loss of migratory capacity in ISR-null cells recovering from protein aggregate stress (6 hrs of 100 nM bortezomib followed by wash). Scratch closure is quantified during an 8 hrs interval, in N=2 independent experiments with n=3-5 technical replicates per condition. Bar graph shows percentage of scratch closure ± SD. *p<0.05 ns, no statistical significance (one-way ANOVA with multiple comparisons).

To directly evaluate focal adhesion dynamics in our cells we generated a zyxin-iRFP construct that would allow for live imaging of cells. Indeed, ISR-null SCC cells recovering from proteotoxic stress possessed large, stable focal adhesions that did not disassemble, rendering the cell immobile over the time frame whereas control SCC cells disassembled their large focal adhesions and displayed considerable dynamics (Figure 6B and Supplemental Video 1). To confirm these aberrant dynamics, we performed a scratch wound assay to induce cell migration following bortezomib treatment. In agreement with our previous data showing large and stable focal adhesion, ISR-null cells showed markedly impaired mobility relative to control cells and failed to migrate into the wound site (Figure 6C and Supplemental Video 2-3). Taken together, these results provided further evidence that in the absence of an ISR, SCC cells exposed to proteotoxic stress are slow to ignite the microtubule dynamics necessary for recovery.

### ATF5 acts downstream of the ISR to promote aggresome formation and the recovery from proteotoxic stress

Our collective data strongly implicated the ISR in regulating microtubule dynamics with the purpose of promoting aggresome assembly and the efficient clearance of protein aggregates that accumulate during proteotoxic stress. Additionally, our analysis of ISR-dependent translational changes, revealed a selective upregulation of centrosomal proteins upon proteotoxic stress (Figure 3G). In particular, we were intrigued by ATF5, which has been described as an integral component of the MTOC (Madarampalli et al., 2015). In agreement with these previous findings, confocal microscopy revealed that upon recovery from proteotoxic stress, ATF5 localized at the centrosome in ISR-competent SCC cells (Figure 7A). Hence, we reasoned that translational upregulation of ATF5 might be needed to regulate microtubule dynamics in the face of proteotoxic stress. To test this hypothesis, we engineered ISR-null cells to overexpress doxycycline-inducible, GFP-tagged ATF5 (Figures 7B and 7C).

**Figure 7:**
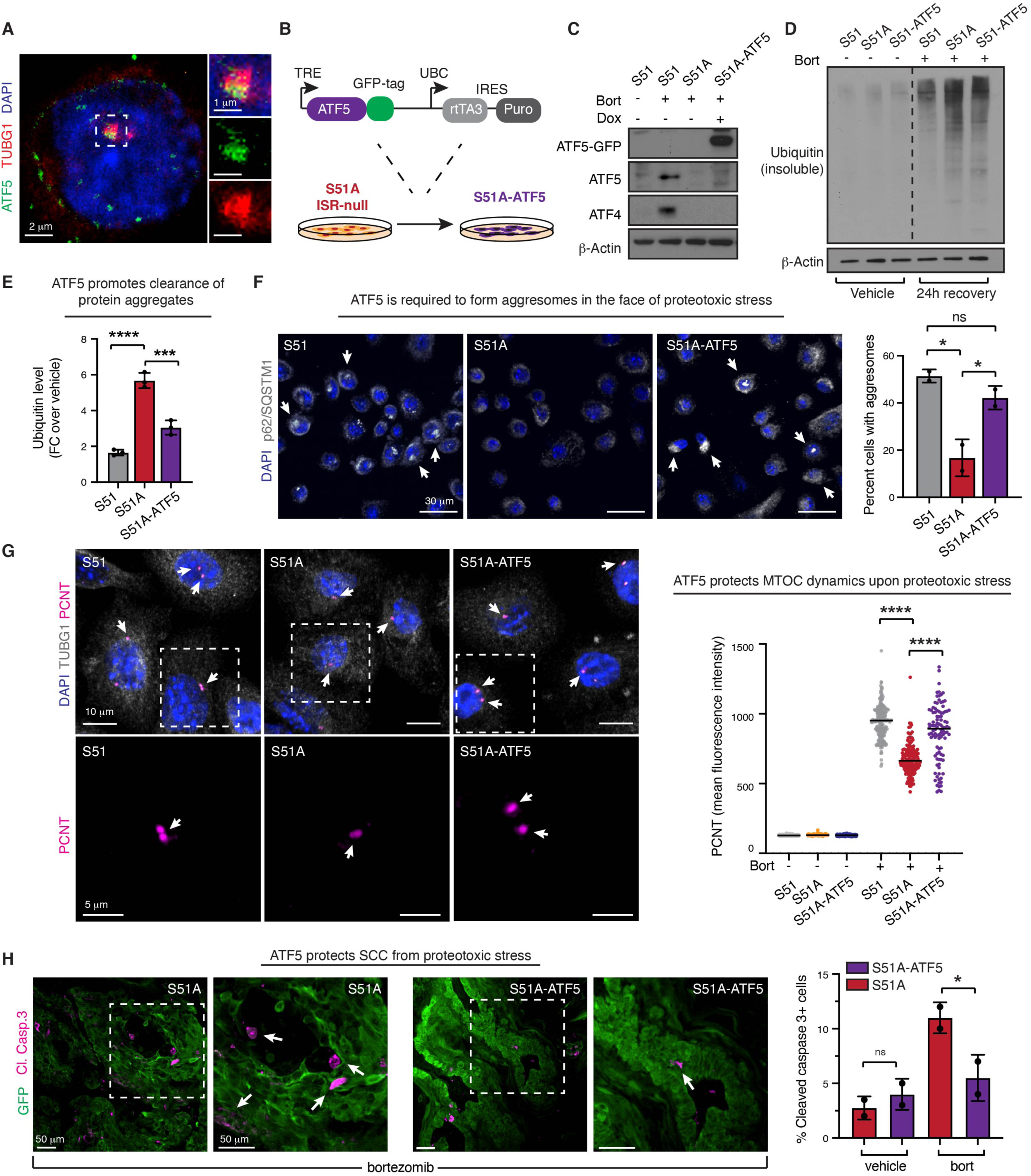
ISR-induced ATF5 protects microtubule dynamics and promotes clearance of protein aggregates. A. Confocal microscopy confirms previously reported ATF5 localization at the MTOC. Insets on the right show higher magnification images with merged (top) and ATF5 (middle) and γ-tubulin (bottom) separately. B. Schematic representation of rescue strategy to express a doxycycline inducible GFP-tagged ATF5 in ISR null SCC cells. C. Immunoblot confirms ATF5 expression upon Dox administration (1 μg/mL for 24 hrs) in proteotoxically stressed S51A-ATF5 cells whose endogenous ATF5 translation is suppressed. D. ATF5 expression partially rescues the ability of proteotoxically stressed ISR-null cells to clear ubiquitinated proteins. Representative immunoblot (of 3 indepdendent biological replicates) shows level of ubiquitinated proteins in the RIPA-insolubule fraction of S51, S51A and S51A-ATF5 SCC cells in the absence or proteotoxic stress or following 24hrs recovery from a 6 hrs bortezomib pulse. E. Quantifications of experiments in (D). Bar graph shows ubiquitin levels ± SD at 24 hrs recovery normalized over actin and relative to ubiquitin levels in untreated cells in 3 independent biological replicates. *p<0.05 ns, no statistical significance (one-way ANOVA with multiple comparisons). F. ATF5 promotes aggresome formation. Aggresomes are visualized by p62/SQSTM1 immunofluorescence in SCC cells treated with a 6 hrs bortezomib pulse followed by a 4 hrs recovery period. White arrows point to aggresomes. Scale bar 30 μm. Bar graph shows percentage of cells with aggresomes ± SD in 2 independent biological replicates. Approximately 300 cells were quantified per each experiment. *p<0.05 ns, no statistical significance (one-way ANOVA with multiple comparisons). G. ATF5 protects centrosomal MTOC dynamics. Centrosomal MTOC is visualized by pericentrin and *γ*-tubulin immunofluorescence. Cells were treated with bortezomib for 6 hrs and let recover for 4 hrs. Scale bar 10 μm. Bottom row shows higher magnification of the area highlighted in top row. White arrows indicate MTOCs. Scale bar 5 μm. Bar graph shows mean fluorescence intensity for pericentrin in 2 independent biological replicates. ****p<0.0001 (one-way ANOVA with multiple comparisons). H. ATF5 induction protects ISR-null cells from protein aggregates-induced cytotoxicity. Immunofluorescence shows SCC tumors (GFP+) after treatment with bortezomib (1.2 mg/kg i.p.). Cell death is quantified by cleaved caspase 3 positive cells. White arrows point to cleaved caspase-3 positive cells. Scale bar 50 μm. Bar graph shows percentage of cleaved caspase 3 positive cells per tumor. *p<0.05 ns, no statistical significance (one-way ANOVA with multiple comparisons).

To understand whether ATF5 is needed to clear protein aggregates, we first examined the clearance of ubiquitinated proteins during the recovery phase following a pulse of bortezomib. Remarkably, expression of ATF5 significantly improved the ability of ISR-null cells to clear ubiquitinated proteins, as assessed by immunoblot of ubiquitinated proteins in the insoluble fraction of cells after 24 hrs recovery from bortezomib (Figure 7D and 7E). Strikingly, this corresponded with a significant increase in aggresome formation, indicating that ISR-mediated ATF5 up-regulation is needed to promote the retrograde transport of misfolded proteins and the subsequent accumulation of protein aggregates at the MTOC (Figure 7F). This effect was mediated via protection of microtubule dynamics as evidenced by increased pericentrin mean fluorescence intensity at the centrosome in ISR-null cells expressing ATF5 after recovery from bortezomib treatment (Figure 7G).

Finally, we asked whether ISR-mediated ATF5 induction promotes cell survival upon proteotoxic stress of SCC cells *in vivo*. To this end, we turned to our grafting model where ISR-null cells displayed increased sensitivity to proteasome inhibition. Consistent with our hypothesis, ATF5 expression rescued the selective sensitivity to bortezomib shown by ISR-null cells. This occurred in the absence of changes in viability in tumors at steady state (Figure 7H). Altogether, these data implicate ATF5 as a critical ISR target, which orchestrates microtubule dynamics to facilitate the clearance of protein aggregates and promote cell recovery upon proteotoxic stress.

## Discussion

Although a handful of reports have linked the actin cytoskeleton to eIF2*α* dephophorylation (Chambers et al., 2015; Chen et al., 2015), cytoskeletal regulation has not been viewed as a primary function of the ISR. In the current study, we unearthed a novel and essential role for the ISR in regulating the cytoskeleton, specifically through preserving microtubule dynamics in stressful situations. We showed that by doing so, proteotoxically stressed cells can accomplish the necessary intracellular trafficking to efficiently clear misfolded proteins and prevent them from accumulating and overtaxing the ER.

Our findings led us to the remarkable conclusion that the ISR not only is required to mount a stress response, but also to maintain cell function during recovery from stress. Although the existence of negative feedback loops that downregulate the ISR upon termination of stress has long been appreciated (Novoa et al., 2001), an active role for the ISR in cellular recovery following a stressful experience has hitherto gone largely unrecognized.

By temporally profiling the translational differences that arise when proteotoxically stressed SCC cells are unable to mount an ISR, we gained insights into the mechanisms underlying the ISR’s importance. Specifically, we learned that when ISR-competent cells are exposed to proteotoxic stress, they redirect their translational machinery to a cohort of mRNAs encoding centrosomal proteins. We show that these proteins function in bolstering the MTOC and reinforcing its microtubule dynamics. This then facilitates efficient microtubule-mediated transport of misfolded proteins to the perinuclear space, where they can be assembled into aggresomes and targeted for ubiquitin-mediated degradation during the stress recovery phase. Indeed, as we showed, when eIF2α cannot be phosphorylated and the ISR core is thereby crippled in proteotoxically stressed cells, the centrosomal proteins are not translated, and microtubule dynamics emanating from the MTOC are disrupted. As judged by deficiencies in the aggresome assembly and in focal adhesion turnover, both retrograde and anterior grade transport of microtubule cargo are slow to recover following stress, thereby impairing cell fitness. Compellingly, the ISR not only upregulates a subset of centrosomal proteins, but also downregulates focal adhesion components, surfacing a network of ISR-regulated cytoskeletal dynamics that directs morphological changes critical for cell recovery.

Microtubule trafficking is important to bring misfolded proteins to the perinuclear space, and our results provided compelling evidence that the ISR functions in this process. However, our findings also pointed to a hitherto unappreciated role of the ISR in aggresome assembly specifically. Our studies revealed that this function is in part mediated by ATF5, a translational target of the ISR in proteotoxically stressed SCC cells and a previously documented structural component of the centrosomal MTOC (Madarampalli et al., 2015). We showed that during recovery from protein aggregate stress, the MTOC size increases concomitantly with aggresome formation, and both of these events depend upon an intact ISR. Our data support a model in which ISR-mediated translational upregulation of centrosomal proteins, including ATF5, is required to remodel the MTOC and concentrate protein aggregates in the perinuclear space so that they can be degraded and cleared by proteosomes and autophagosomes.

Several studies have provided tantalizing evidence that some cells may be able to asymmetrically partition their centrosomal aggregates when division resumes following proteotoxic stress, such that one daughter (e.g. a stem cell) remains healthy, while the other daughter inherits the aggregates and becomes slated to differentiate (Morrow et al., 2020; Rujano et al., 2006; Singhvi & Garriga, 2009). We did not see evidence for this in our SCC cells. Rather, there was a lag in cell cycle reentry such that it coincided with the timing at which accumulated ubiquitinated misfolded proteins were cleared (Figures 2D and 5B). This coupling was especially striking in comparing the behaviors of ISR-null versus ISR-competent cells. Thus, during the recovery phase, ISR-null cells were delayed by > 24 hrs in both aggregate clearance and cell cycle re-entry. That said, when the ISR was crippled either *in vivo* or *in vitro*, SCC cells were jeopardized in their overall ability to survive proteotoxic stress. In this scenario, as a major regulator of cell survival in response to proteotoxic stress, our findings suggest that by coupling pharmacologic inhibition of the ISR (Sidrauski et al., 2015) with proteosomal inhibitors such as bortezomib, such a regimen may find an Achilles heel for this family of difficult to treat cancers.

### Ideas and speculation

Our findings are also likely to have relevance for other diseases, including neurodegenerative diseases, which are driven by or associated with protein aggregates (Soto & Pritzkow, 2018). To date, the function of the ISR in neurodegenerative diseases has been attributed largely to the emergence of alternative translation pathways and to ATF4 transcriptional activity that drives expression of cytoplasmic chaperones and balances cell survival and cell death (Bond et al., 2020; Costa-Mattioli & Walter, 2020). In SCC cells, although it is possible that chaperones may be upregulated indirectly through ISR-driven transcription factors, these proteins did not emerge as major translational targets of the ISR in our ribosome profiling analysis. Moreover, since cells have parallel pathways, most notably the heat shock response, that can upregulate chaperones when needed (San Gil et al., 2017), it seems unlikely that this would be ISR’s sole function in maintaining proteostasis.

The necessity of an ISR-driven pathway to preserve centrosome dynamics during proteotoxic stress raises an important question regarding centrosome function in neurodegenerative disorders. If ISR-driven translation of centrosomal proteins is required to protect microtubule dynamics during stress, it follows that the accumulation of unfolded proteins should negatively impact the MTOC and/or its microtubule-associated dynamics. In Alzheimer’s disease and other tauopathies, the microtubule associated protein Tau is the main driver of the aggregates of neurofibrillary tangles that ensue, and this alone is likely to negatively impact microtubule-mediated trafficking in neurons (Ballatore et al., 2007). In addition, misfolded protein aggregates often inadvertently sequester properly folded cellular proteins, such as p62 and the disaggregase p97/VCP, which can exacerbate their toxic effects on cells (Donaldson et al., 2003; Olzscha et al., 2011; Yang & Hu, 2016). It seems plausible that misfolded protein aggregates might either sequester low-complexity centrosomal or pericentrosomal proteins (Woodruff et al., 2017), or structurally interfere with MTOC function. In fact, one study found that proteotoxic stress does indeed inhibit centrosome function in neurons (Didier et al., 2008), raising the question as to whether the ISR might also be involved in maintaining the MTOC of long-lived, non-dividing neurons, which face challenging microtubule-mediated cellular trafficking dynamics of the likes that few if any other cell type of the body does.

In summary, our work supports a model in which most if not all cells, but likely cancer cells and neurons in particular, rely upon the ISR to restore proteostasis following protein aggregate stress. By redirecting the translational machinery towards synthesizing proteins involved in enlarging the MTOC and bolstering microtubule dynamics, the ISR aids in the intracellular microtubule-mediated trafficking necessary to assemble aggresomes at the MTOC, where they can be efficiently targeted to the perinuclear protein degradation machinery.

## Materials and methods

### CRISPR cloning

Lentiviral particles containing the gene replacement construct, pLKO-PGK-eIF2α-mycTag-P2A-NeoR were prepared by transfecting the lentiviral plasmid along with packaging plasmids into HEK293T cells, and viral supernatant was collected 48 hours post transfection. This construct was integrated into the genome of a clonal, parental primary SCC mouse line by incubating 100 μL with 0.1 mg/mL polybrene for 8 hours. 48 hours later cells with the integrated construct were selected with 0.5 mg/mL Neomycin selection for 2 days. Following selection the endogenous allele was targeted for deletion using CRISPR-Cas9 RNP particles. The replacement allele had a synonymous mutation in the PAM site rendering it resistant to this CRIPSR construct. CRISPR-Cas9 RNP particles targeting the endogenous allele were prepared as follows: eIF2α gRNA (target sequence: ATATTCCAACAAGCTGACAT) was designed using Guidescan software (Perez et al., 2017) and complexed with ATTO550-tracrRNA and Cas9. All reagents were acquired from IDTdna’s “AltR system”. Duplexed gRNA:tracrRNA was prepared by mixing 1 μM of each component in IDTdna duplex buffer, heated to 95° C in a thermocycler and annealed by gradually lowering the temperature to 25° C at a rate of 0.1° C/second. Duplexed gRNA:tracrRNA was complexed with Cas9 by mixing 1 μM of the duplex with 1 μM Cas9 in OptiMEM (ThermoFisher) and incubating at room temperature for 5 minutes. RNP complexes then were transfected into 60% confluent 12 well plates using RNAiMAX as follows: 30 μL of 1 μM RNP complexes were mixed with 4.8 μL RNAiMAX in 335 μL of optiMEM and RNP-lipid complexes were allowed to form for 15 minutes at room temperature. At the end of the incubation 400 μL of complexes were added dropwise to cells, and media was changed 16 hours later. 48 hours after transfection single cells were isolated using fluorescence activated cell sorting (FACS) as follows: Cells were dissociated with trypsin (Gibco) and resuspended in 500 μL of FACS buffer (PBS supplemented with 5% FBS and 5 uM EDTA), and single, ATTO-550 positive cells were sorted using BD FACSAria cell sorter into wells of 96-well plates containing 100 μL of 50-50 mixture of fresh media and conditioned media. Clones that grew to confluency were transferred to 12 well plates, and following growth to confluency in 12 well plates, cells were dissociated with trypsin and frozen in freezing media supplemented with 10% FBS and 10% DMSO. At this stage a small aliquot of cells (75% of the plate) was lysed in 200 μL of QuickExtract DNA Extraction solution (Lucigen) and gDNA was prepared by heating to 65° C for 10 minutes followed by heat inactivation at 95° C for 2 minutes. These gDNA samples were used for further analysis. For HRI KO cells the protocol was exactly the same except that gRNA targeting HRI (target seq: ATTTAAACACCTGTTTGGAG) was used.

### NGS analysis of CRISPR outcomes

Knockout of the endogenous allele was evaluated using primers targeting a 300 basepair region of genomic DNA with the targeted locus in the middle of the amplicon. Primers had 5’ overhangs with sequences compatible with the Illumina Nextera XT index primers (R: overhang: GTCTCGTGGGCTCGGAGATGTGTATAAGAGACAG, L overhang: TCGTCGGCAGCGTCAGATGTGTATAAGAGACAG). Amplicons were generated with 1 μL of input gDNA and the NEB phusion kit according to manufacturer’s instructions, and amplicons were isolated using Agencout Ampure XP beads (Beckman Coulter). A second barcoding PCR was performed using Nextera XT index primers as follows: 2 μL of cleaned amplicons were used as input, primers were added so that each isolated clone had a unique combination of left and right barcodes, and barcodes were added using a 8 cycle PCR reaction with 55° C annealing temp, again with the NEB Phusion kit, and the barcoded amplicons were cleaned and primer dimers were removed using Ampure XP beads. Amplicons were normalized to the same concentration, pooled, and sequenced using a single Illumina MiSeq Nano lane using the 250 basepair, paired end kit. Demultiplexed reads were analyzed and screened for indels using the RGEN Cas Analyzer (http://www.rgenome.net). A KO clone was confirmed if the only reads detected in that sample were indels that would create a frameshift.

### Cell culture

Primary murine SCC cells were generated and cultured in E medium supplemented with 15% FBS and 50 mM CaCl2 as previously described (Yang et al., 2015). Cells were passaged 3 times per week and passage numbers were maintained counting from the point of cell line generation. Frozen cell stocks were generated by freezing cells in complete media supplemented with 10% additional FBS and 10% DMSO. ATF5-GFP overexpressing cells were generated by infecting ISR-null cells with ATF5-GFP lentivirus in the presence of 5 μg/mL polybrene. Infected cells were selected using 2 μg/mL puromycin for 5 days. Induction of ATF5 was induced using 1 μg/mL doxycycline at the time of plating for eachexperiment.

### Proliferation and cell viability assays

For proliferation assays 2500-5000 cells were plated per well in clear-bottom, black optical 96 well plates (Nunc) and allowed to attach overnight. A baseline plate was collected the next day as a zero hour sample, and then plates were collected at 24, 48, and 72 hours after this timepoint. At the time of collection media was washed and cells were fixed in 4% PFA in PBS for 10 minutes at room temperature. After fixation, PFA was washed with PBS and cells were stored in PBS at 4° C until the end of the experiment. Following collection of the final plate, nuclei were stained in all samples using 1 ug/mL DAPI in PBS for 5 minutes at room temperature. DAPI was washed and replaced with PBS, and nuclei were imaged on a Biotek Cytation 5 high content imager, and cells were counted using Gen5 software. To measure cell death, single-cell-suspensions were washed once with PBS, stained with 3 μl of AnnexinV-PE conjugates (Thermo Fisher) and 0.1 μg/ml DAPI (Thermo Fisher) in 100 μl of 1X Annexin Binding Buffer (Thermo Fisher) for 15 min at room temperature. Data were collected using a BD LSR Fortessa X20 flow cytometer and analyzed using FlowJo software.

### Measurement of translation rates

Cells were plated to 75% confluency in 6 well plates. The next day cells were treated with 50 μM sodium arsenite, 100 ng/mL tunicamycin or 100 nM borteozomib, or a vehicle control and incubated at 37° C for 6 hrs. At this time 20 μM puromycin was supplemented to the media, and cells were incubated for 30 min. Cells were then lysed in RIPA buffer and translation rates were evaluated by immunoblotting for puromycilated peptides using an anti-puromycin antibody.

### Microtubule nucleation assay

Cells were plated to 50% confluency in glass slides (Millicell EZ, Millipore Sigma) and treated with 100 nM bortezomib or a vehicle control for 6 hrs followed by a PBS wash and replacement with fresh media. 4 hrs later the media was supplemented with 13 μM nocodazole, and cells were incubated at 37° C for 20 mins. At the end of the incubation slides were washed and media was replaced with fresh, nocodazole-free media, and microtubules were allowed to recover during a 2 min incubation at room temperature. At this time cells were fixed in 4% PFA for 10 min at room temperature. Additional control slides without nocodazole or with nocodazole and no recovery were fixed as controls. Slides were processed for immunofluorescence targeting α-tubulin, pericentrin, and λ-tubulin, immunofluorescence signal was imaged by confocal microscopy, and microtubule nucleation rates were evaluated by quantifying the relative α-tubulin signal intensity within the centrosome region, which was defined using 3D volumetric assessment (Imaris) of pericentrin-positive volumes.

### Scratch assay

Cells were plated in plastic bottom optical slides (Ibidi 80826) and allowed to reach confluency in in 48 hours. At this time cells were treated with 100 nM bortezomib or a vehicle control for 6 hours followed by PBS wash and replacement with fresh, drug-free media. At the time of wash scratch wounds were manually created by gently scaping cells using a rubber cell scraper. Movement of wound-edge SCC cells was evaluated using time-lapse confocal microscopy of GFP signal (marking all cells) with images acquired every 2 minutes for 8 hours. Scratch closure was evaluated using Imaris software to measure the percentage of scratch closed by leading edge cells over the course of the experiment.

### FACS isolation of ex vivo tumor cells

To sort SCC cells out of tumors formed from ISR-WT and ISR-null cells lines, Day 35 tumors were dissected from the skin and finely minced in 0.25% of collagenase (Sigma) in HBSS (Gibco) solution. The tissue pieces were incubated at 37° C for 20 minutes with gently shaking. After a single wash with ice-cold PBS and samples were further digested into single cell suspension in 0.25% Trypsin/EDTA (Gibco) for 10 min at 37° C. After neutralization with the FACS buffer (PBS supplemented with 4% FBS, 5 mM EDTA, and 1 mM HEPES), single-cell suspensions was then centrifuged, resuspended, and strained before preparing for staining. A cocktail of Abs for surface markers at the predetermined concentrations (CD31-APC 1:100, Biolegend; CD45-APC 1:200, Biolegend, CD117-APC 1:100, Biolegend; CD140a-APC 1:100, Thermo Fisher; CD29-APCe780, 1:250, Thermo Fisher, Biolegend) was prepared in the FACS buffer with 100 ng/ml DAPI. Sorting was performed using a BD FACSAria equipped with FACSDiva software to isolate a population of cells that was GFP-positive (pan-SCC), DAPI-negative (live), and APC-negative (dump gate to exclude immune cells, endothelial cells, and fibroblasts), and new cell lines were established from bulk populations of sorted cells.

### Tumor allografting

Squamous cell carcinoma allografts were generated by intradermally injecting 1×10^5^ SCC cells suspended in a 50:50 mix of PBS and growth-factor reduced Matrigel (Corning, 356231) in an injection volume of 50 μL. Grafts were generated in the flanks of 6-8 week old female (*Nude*) mice. Tumor dimensions were measured every 5 days using electronic calipers and tumor volume was calculated using the formula V = 0.5 × length × width2. For tumor growth experiments sample sizes of N=8 were calculated to yield an 80% power to detect a significant (p<0.05) effect size of 50% assuming standard deviation of 25%, conservate estimates based on past lab experience suggesting targeting ISR-related genes in SCC which yielded larger effect sizes (Sendoel et al., 2017). Experiment was performed twice ISR-WT and ISR-null using two different clones for each, with each experiment powered independently to 80%, yielding total sample size of N=16 when pooling data. Experiments evaluating apoptosis *in vivo* were set up for 80% power to detect a significant (p<0.05) effect size of 66% with 25% SD, which yielded minimum sample size of N=4.

### Immunofluorescence/histology

For immunofluorescence of tumors, samples were fixed in 4% PFA for 1 hr at room temperature , dehydrated in 30% sucrose overnight at 4°C, and mounted into OCT blocks and frozen. 14 μm thick sections were cut using a Leica cryostat deposited onto SuperFrost Plus slides (VWR). For immunofluorescence of cells in culture, cells were plated on glass slides (Millicell EZ, Millipore Sigma) coated with human plasma fibronectin (Millipore Sigma) diluted to 100 μg/mL in PBS. At the conclusion of experiments samples were fixed with 4% PFA for 10 minutes at room temperature. Samples were permeabilized with 0.3% Triton-X100 in PBS and blocked using 2.5% normal donkey serum, 2.5% normal goat serum, 1% BSA, 2% fish gelatin, and 0.3% Triton X-100 in PBS. Primary antibodies were applied in blocking buffer overnight at 4° C. Samples were washed with 0.1% Triton X-100 and secondary antibodies with Alexa 488, Alexa 594, and Alexa 647 were applied for 1 hour at room temperature in blocking buffer containing 1 μg/mL DAPI. Slides were washed with 0.1% triton and mounted using Prolong Diamond Antifade Mountant with DAPI (ThermoFisher).

### Microscopy and image analysis

Microscopy of tumors and 40X images of aggresomes and spreading cells were performed using an Axio Observer Z1 epifluorescence microscope equipped with a Hamamatsu ORCA-ER camera (Hamamatsu Photonics), and with an ApoTome.2 (Carl Zeiss) slider using a 20X air, 40X oil, or 63X oil objective. 63X confocal microscopy images were collected on an Andor Dragonfly spinning disk imaging system with a Leica DMi8 Stand and cMOS Zyla camera. Images were analyzed in FIJI or Imaris. For fluorescence intensity measurements of tumors GFP masks were generated and signal was measured within the mask. Aggresomes were manually counted as discrete p62-positive juxtanuclear puncta on maximum intensity Z-projections. Cell dimensions were calculated using the length measuring tool, and cell spreading was evaluated manually by observing for spread morphology. RGB images were generated with FIJI and saved as TIFF files. For 3-dimensional reconstructions and volumetric analyses of microtubule organizing centers, Imaris was used to generate 3D images, and volumes were generated to create 3D volumes encompassing the discrete puncta of pericentrin staining. Volume as well as summed pericentrin and γ-tubulin fluorescence intensity were measured within these volumes

### Electron microscopy

Cells were fixed in a solution containing 4% PFA, 2% glutaraldehyde, and 2 mM CaCl2 in 0.1 M sodium cacadylate buffer (pH 7.2) for 1 hr at room temperature, and then placed at 4° C. Cells were next postfixed in 1% osium tetroxide and processed for Epon embedding; ultrathin sections (60-65 nm) were then counterstained with uranyl acetate and lead citrate, and images were acquired using a Tacnai G2-12 transmission electron microscope equipped with an AMT BioSprint29 digital camera.

### Cell fractionation

Cells were lysed in RIPA buffer (20 mM Tris-HCl, pH 8.0, 150 mM NaCl, 1 mM EDTA, 1 mM EGTA, 1% Triton X-100, 0.5% deoxycorate, 0.1% SDS) containing protease inhibitors (Complete mini, Roche) and phosphatase inhibitors (PhosStop), and lysed for 10 minutes at room temperature. Membrane fraction was pelleted by centrifuging at 1000 g for 10 minutes at 4° C. The supernatant (cytosolic fraction) was transferred to new tubes which were centrifuged at 20,000 g for 30 min at 4° C. The supernatant (RIPA-soluble fraction was transferred to a new tube, and the pellets (RIPA-insoluble fractions) were washed with 300 μL of RIPA buffer, centrifuged 20,000 g for 10 min at 4° C, and then resuspended in 30 μL of 1X LDS-βME by vortexing vigorously and boiling at 98° C for 10 minutes. The protein concentration of the RIPA-soluble fraction was measured with a BCA assay (Pierce). RIPA-soluble fractions were mixed into 1X LDS-βME and protein concentration was normalized. The insoluble fractions were normalized by adding 1X LDS-*β*Me so that the same volume corresponds to the same volume of insoluble-fraction lysate (eg. Insoluble fraction from 100 μg of cell lysate). Samples were run on immunoblots as previously described and probed for ubiquitin signal.

### Western blotting

Cells were lysed with RIPA (20 mM Tris-HCl, pH 8.0, 150 mM NaCl, 1 mM EDTA, 1 mM EGTA, 1% Triton X-100, 0.5% deoxycorate, 0.1% SDS) containing protease inhibitors (Complete mini, Roche) and phosphatase inhibitors (PhosStop). Lysates were clarified by centrifugation and protein concentration of supernatants was evaluated using BCA assay (Pierce). Protein lysates were normalized, mixed with LDS (ThermoFisher) and β-mercaptoethanol (Thermofisher), and denatured at 95° C for 5 minutes. Protein was separated by gel electrophoresis using 4–12% or 12% NuPAGE Bis-Tris gradient gels (Life Technologies) and transferred to nitrocellulose membranes (GE Healthcare, 0.45 μm). Membranes were blocked with 2% BSA in TBS supplemented with 0.1% Tween 20, and primary antibodies were stained overnight in blocking buffer at 4° C or overnight at room temperature, and HRP-conjugated secondary antibodies were stained for 1 hour at room temperature. Membranes were washed and then incubated with ECL plus chemiluminescent reagent (Pierce) for 30 seconds. Chemiluminescent signal was evaluated using CL-XPosure Film (ThermoFisher).

### Antibodies and counterstains

The following antibodies and dilutions were used for immunoblotting. 1/1000 β-Actin (Cell Signaling Technologies, 3700); 1/1000 p-eIF2α (Invitrogen, 44-728G); 1/5000 p-eIF2α (Cell Signaling Technologies, 3597); 1/1000 eif2a (Cell Signaling Technologies, 9722) 1/3000 puromycin (Millipore Simga MABE343); 1/1000 p62/SQSTM1 (Cell Signaling Technologies, 5114); 1/1000 Atf4 (Cell Signaling Technologies, 11815); 1/1000 ubiquitin (Cell Signaling Technologies, 3393S); 1/1000 K48-linked polyubiquitin (Cell Signaling Technologies, 8081); 1/1000 Myc-tag (Abcam, ab32); 1/5000 α-tubulin (Millipore Sigma, T5168); and 1/1000 Atf5 (Santa Cruz, sc377168). For immunofluorescence the following antibodies and dilutions were used. 1/500 α6-integrin (BD Bioscience, 555734); 1/500 GFP (Abcam, ab13970); 1/500 E-Cadherin (Cell Signaling Technologies, 3195); 1/100 G3BP (BD Bioscience 611126); 1/500 Vinculin (Millipore Sigma, V9131); 1/100 Phospho-Myosin Light Chain 2 (Ser19) (Cell Signaling Technologies, 3675); 1/250 p62/SQSTM (Cell Signaling Technologies, 7695); 1/500 LaminB1 (Santa Cruz, 374015); 1/500 α-tubulin (BD Biosciences, MCA77G); 1/500 γ-tubulin (Millipore Sigma, T6557); 1/250 Pericentrin (Abcam, ab4448) 1/200 Atf5 (Abcam ab184923). Nuclei were counterstained with 1 μg/mL DAPI (ThermoFisher) and F-Actin was stained with 1/400 Rhodamine or AlexaFluor-488-conjugated phalloidin (ThermoFisher).

### Ribosome profiling and analysis

To perform ribosome profiling we closely followed a recently published protocol (McGlincy & Ingolia, 2017). In short cells were lysed in polysome buffer supplemented with 0.1 mg/mL cyclohexamide. Lysates were treated with 500 U of RNAse I (Epicentre) per 25 μg of RNA (quantified by Qubit fluorimetry – ThermoFisher), and ribosome protected fragments were isolated using sephacryl S400 columns (GE Healthcare) in TE buffer. Ribosomes and ribosome protected fragments (RPFs) were dissociated in Trizol (Invitrogen) and RNA was collected with protected fragments were purified using Zymo Research DirectZol MiniPrep columns. RPFs were purified by running total RNA on a 15 % TBE-Urea gel (ThermoFisher) and cutting the region corresponding to 17 to 34 nucleotides. RNA was purified and precipitated, dephosphorylated with PNK (NEB), and ligated to preadenylated barcoded linker oligonucleotides, and then unligated linker was digested away with 25 U Yeast 5’-deadenylase (NEB) and 5 U RecJ exonuclease (Epicentre). Up to 8 libraries were pooled, and rRNA was depleted using the Lexogen Ribocop V2 kit, and reverse transcription of RPF-linker fragments was performed in presence of 20U Superase-IN (Invitrogen) and 200U Protoscript II (NEB). cDNAs were circularized using 100U CircLigase I ssDNA ligase (Epicentre), and cDNA concentration was quantified by QPCR of cDNA compared to a standard curve of a reference sample. Libraries were amplified with Illumina-compatible barcoded primers and a 10 cycle PCR. DNA of the correct size (∼160 bp) was isolated on TBE-PAGE gel, precipitated, and resuspended in TE buffer. Total mRNA samples were prepared in parallel, and mRNA was selected by rRNA depletion, and libraries were prepared using Illumina Ribozero kit.

Ribosome profiling and total RNA libraries were pooled and sequenced on a Novaseq using the S1, 1×100 bp kit. Reads were demultiplexed, trimmed using FastX trimmer, and aligned to the mm10 reference genome using bowtie2. Sequences were counted in bins of 5’ UTR, CDS, and 3’ UTR as defined using plastid (https://plastid.readthedocs.io/en/latest/). Count data was analyzed using DESeq2 (Love et al., 2014), an R package designed for statistical analysis of gene counts generated from Illumina-based sequencing. For expression analysis of ribosome profiling data only reads in the CDS were included and reads coming from the first 15 codons (45bp) and last 5 codons (15bp) were excluded. Additionally, only reads of size between 20-23bp or 26-32bp were counted. Gene lists were generated as described in the main text, first by filtering genes with significant differences in RPF-read counts, and then genes that changed specifically at the translation level were identified as the subset of filtered genes with translational efficiency (TE=normalized RPF/normalized total RNA) fold changes greater than 1.5.

## Data availability

The total mRNA and ribosome protected fragments datasets generated during the ribosome profiling experimetns performed in this study have been deposited to the Gene Expression Omnibus (GEO) repository with accession code GSE193945.

## ACKNOWLEDGMENTS

We thank Rockefeller University’s Flow Cytometry Resource Center (Svetlana Mazel, director), the Genomics Resource Center (Connie Zhao, director) and the American Association for the Accreditation of Laboratory Animal Care-accredited comparative biology center (R. Tolwani, director) for their services. We thank the Epigenomics Core Facility (Yushan Li, director) at Weill Cornell University for sample processing. We also thank members of the Fuchs lab, specifically Lisa Polak for assisting with the engraftments and tumor studies, and Dr. Stephanie Ellis and Dr. Katherine S. Stewart for their helpful discussions and for assistance in setting up various experiments performed in this study. B. H. is a Ruth Kirschstein NIH Predoctoral Fellow (F30CA236239) and a member of the Weill Cornell/Rockefeller/Sloan Kettering Tri-Institutional Medical Scientist Training Program (T32GM007739). N.G. is a HHMI Jane Coffin Childs Associate. A.G. is a Damon Runyon Cancer Research Foundation National Mah Jongg League Fellowship (DRG 2409-20). E.F. is an investigator of the Howard Hughes Medical Institute. The work is also supported by grants from the National Institutes of Health (R01-AR27883 to E.F.) and the Robertson Foundation.

---

**Figure 1-Figure supplement 1:**
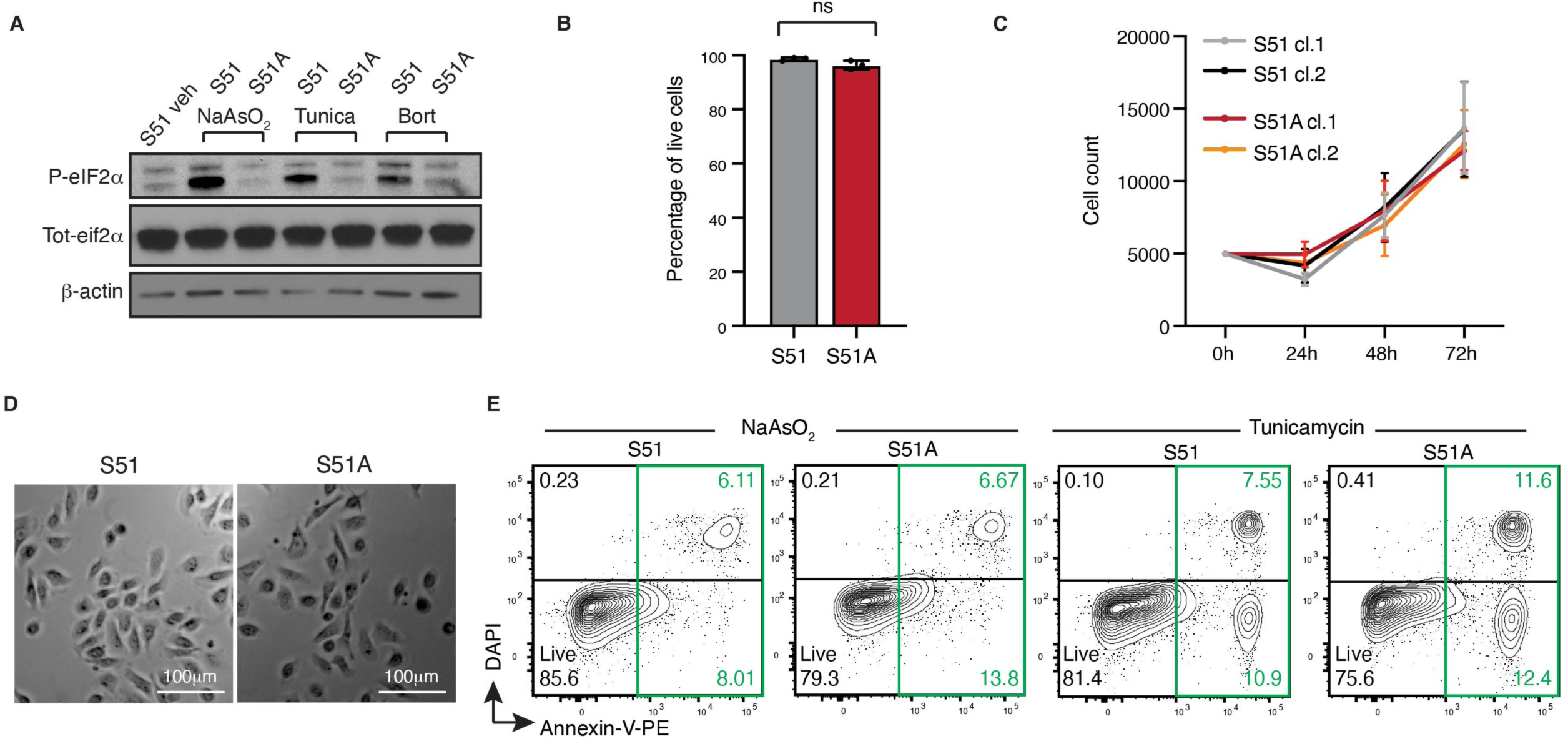
ISR-null cells are more sensitive to stress, but show no difference at steady state. A. ISR-competent cells induce eIF2*α* phosphorylation upon stress. Representative immunoblot shows levels of eif2*α* phosphorylation of cells treated with either 50 μM sodium arsenite, 100 ng/mL tunicamycin or 100 nM bortezomib for 6 hrs. B. Bar graph shows no difference in viability in control and ISR-null cells at steady state. ns, no statistical significance (t test). C. Quantification of cell count over time demonstrates no difference in proliferation rate between ISR-competent and ISR-null cells. Representation of N=4 experiments and n=16 total technical replicates per condition, error bars denote SD of pooled technical replicates. D. Representative brightfield microscopy images of control and ISR-null cells show similar morphology in the absence of stress. D. ISR-null cells are more sensitive to stress. Representative FACS plot show increased annexin-V staining in ISR-null cells after 24 hrs treatment with 5 μM sodium arsenite or 100 ng/mL tunicamycin.

**Figure 2-Figure supplement 1:**
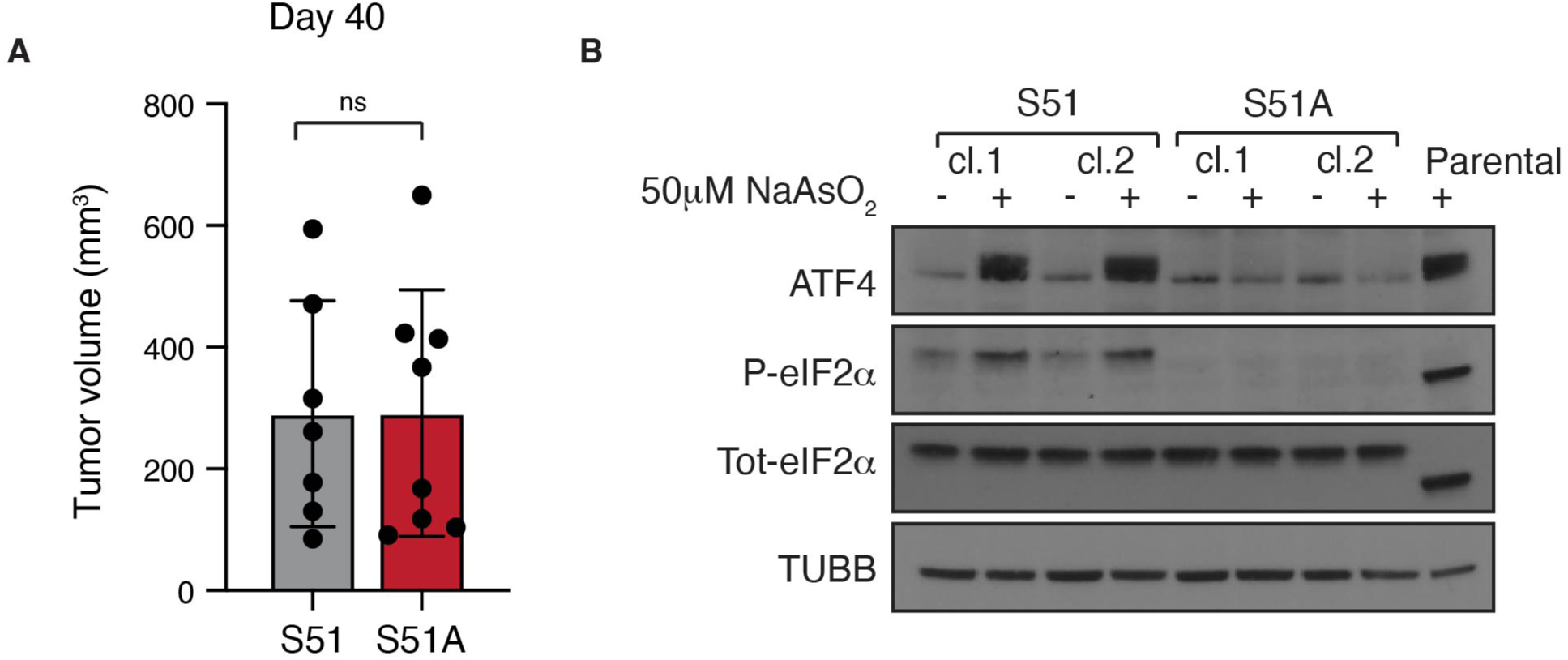
Characterization of ISR-null SCC tumors. A. Bar graph shows tumor volume quantification ± SD at day 40 in mice transplanted with either control or ISR-null SCCs. A minimum of 7 tumors per condition were quantified. ns, no statistical difference (t test). B. Secondary ex vivo control and ISR-null SCC cell lines were generated by FACS isolation of day 35 tumors initiated and grown from transplantation of our primary control and ISR-null SCC lines. p-eIF2α and ATF4 immunoblot following 4 hrs treatment with 50 μM sodium arsenite shows that neither ISR-null clone (N=2) has escaped ISR-ablation. This experiment rules out the trivial explanation that ISR-null SCC growth is due to ‘escaper’ cells that have reverted to a wild-type ISR state.

**Figure 2-Figure supplement 2:**
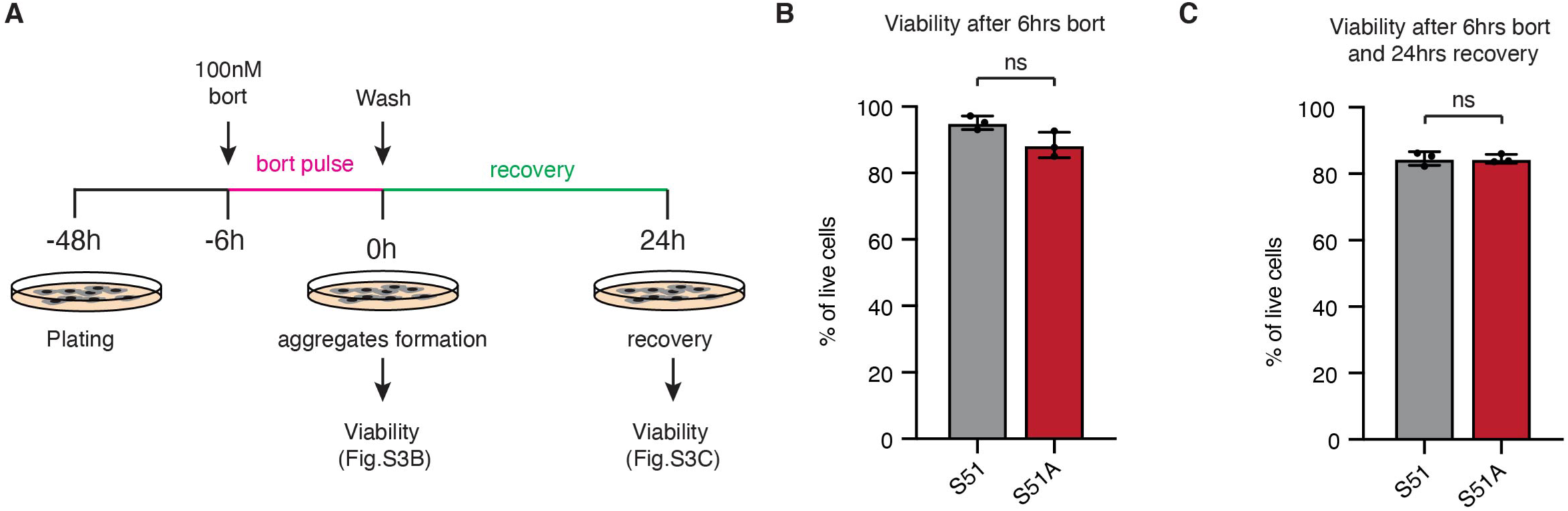
Characterization of bortezomib pulse and recovery system. A. Schematic illustrates experimental design for our bortezomib pulse and recovery system. Cells are treated for 6 hrs with 100 nM bortezomib and let recover for 24 hrs. Viability is assessed at two time points (0 hrs recovery and 24 hrs recovery) B. ISR-null cells show no difference in viability after a short (6 hrs) pulse of 100 nM bortezomib. Bar graph shows percentage of live cells ± SD quantified as DAPI negative cells by FACS in three independent experiments. ns, no statistical significance (t test). C. ISR-null cells show no difference in viability after a 24 hrs recovery from a 6 hrs pulse of 100 nM bortezomib. Bar graph shows percentage of live cells ± SD quantified as DAPI negative cells by FACS in three independent experiments. ns, no statistical significance (t test).

**Figure 2-Figure supplement 3:**
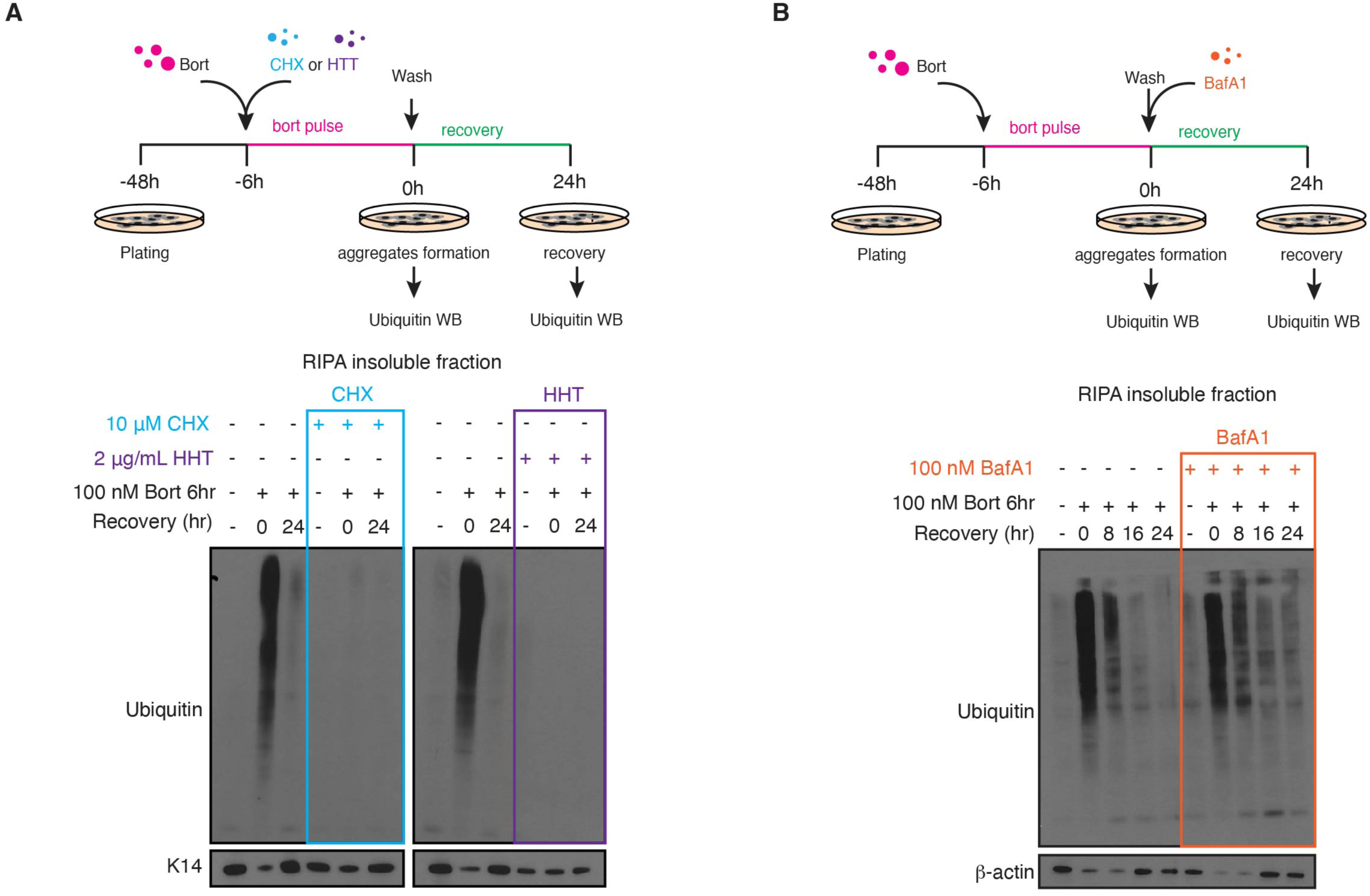
Characterization of protein aggregate source and clearance mechanism. A. Protein aggregates formation is dependent on newly synthesized proteins. Ubiquitin immunoblot in the RIPA-insoluble fraction shows absence of protein aggregates in control cells either treated with 10μM of the translation elongation inhibitor cycloheximide or 2 μg/mL of the translation initiation inhibitor harringtonine concomitantly with the bortezomib pulse. B. Autophagy is only partially required to clear protein aggregates. Ubiquitin immunoblot in the RIPA-insoluble fraction shows partial accumulation of protein aggregates in presence of 100 nM of the autophagy inhibitor bafilomycin A1.

**Figure 3-Figure supplement 1:**
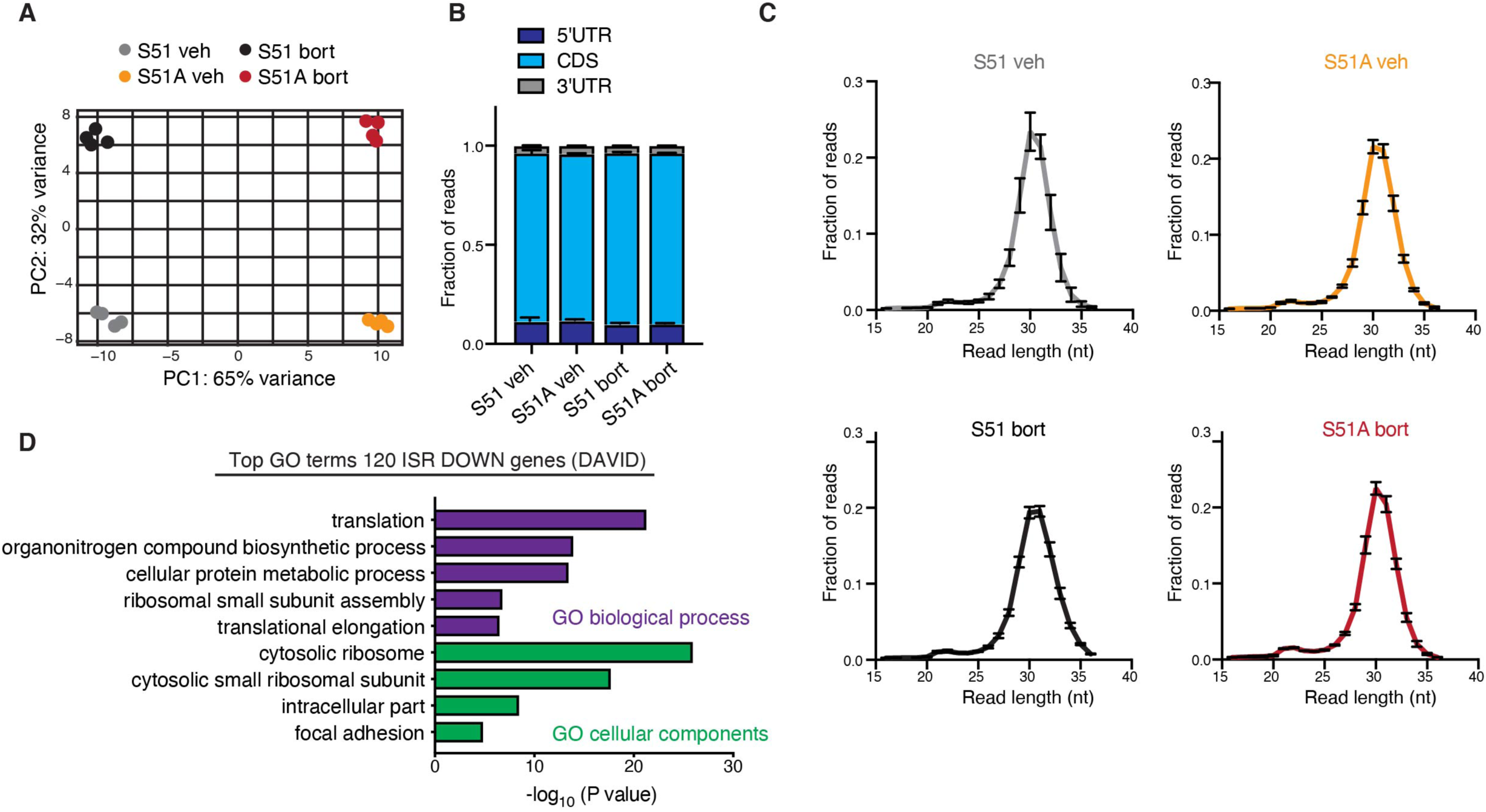
Quality control of ribosome profiling libraries. A. PCA plot shows variance of 4 independent replicates. Two major sources of variance are identified in our dataset: S51 vs S51A (x-axis) and vehicle vs bortezomib (y-axis). B. Metagene analysis shows fraction of RPF reads respectively aligned to 5’UTR, CDS and 3’UTR. C. Histograms show fraction of RPF reads corresponding to their length in nucleotides. As expected the majority of reads are found between 29-30 nt. D. GO analysis shows top terms for biological processes and cellular components of 120 genes translationally downregulated by the ISR. 120 genes are defined by the overlap of mRNAs with significantly downregulated translation in two comparisons: (1) ISR-competent-bort vs. ISR-competent-vehicle and (2) ISR-competent-bort vs. ISR-null-bort.

**Figure 4-Figure supplement 1:**
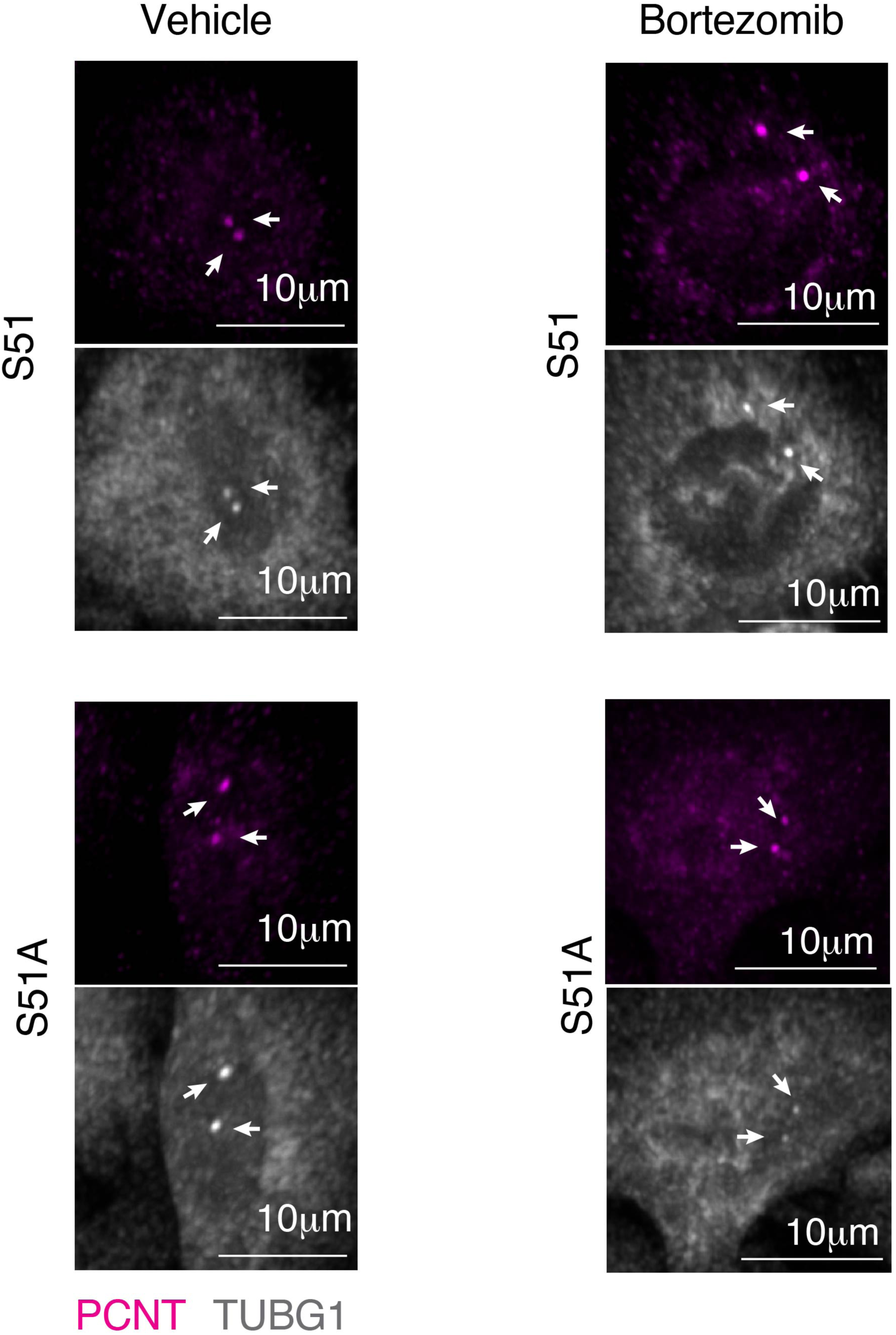
Increased centrosomal MTOC size upon proteotoxic stress is dependent on the ISR. High magnification images show centrosomal MTOC by pericentrin and *γ*-tubulin staining in S51 and S51A cells vehicle-treated or treated with a bortezomib pulse (100 nM, 6 hrs) and let to recover for 4 hrs. White arrows indicate MTOCs. Scale bar 10 μm.

**Figure 5-Figure supplement 1:**
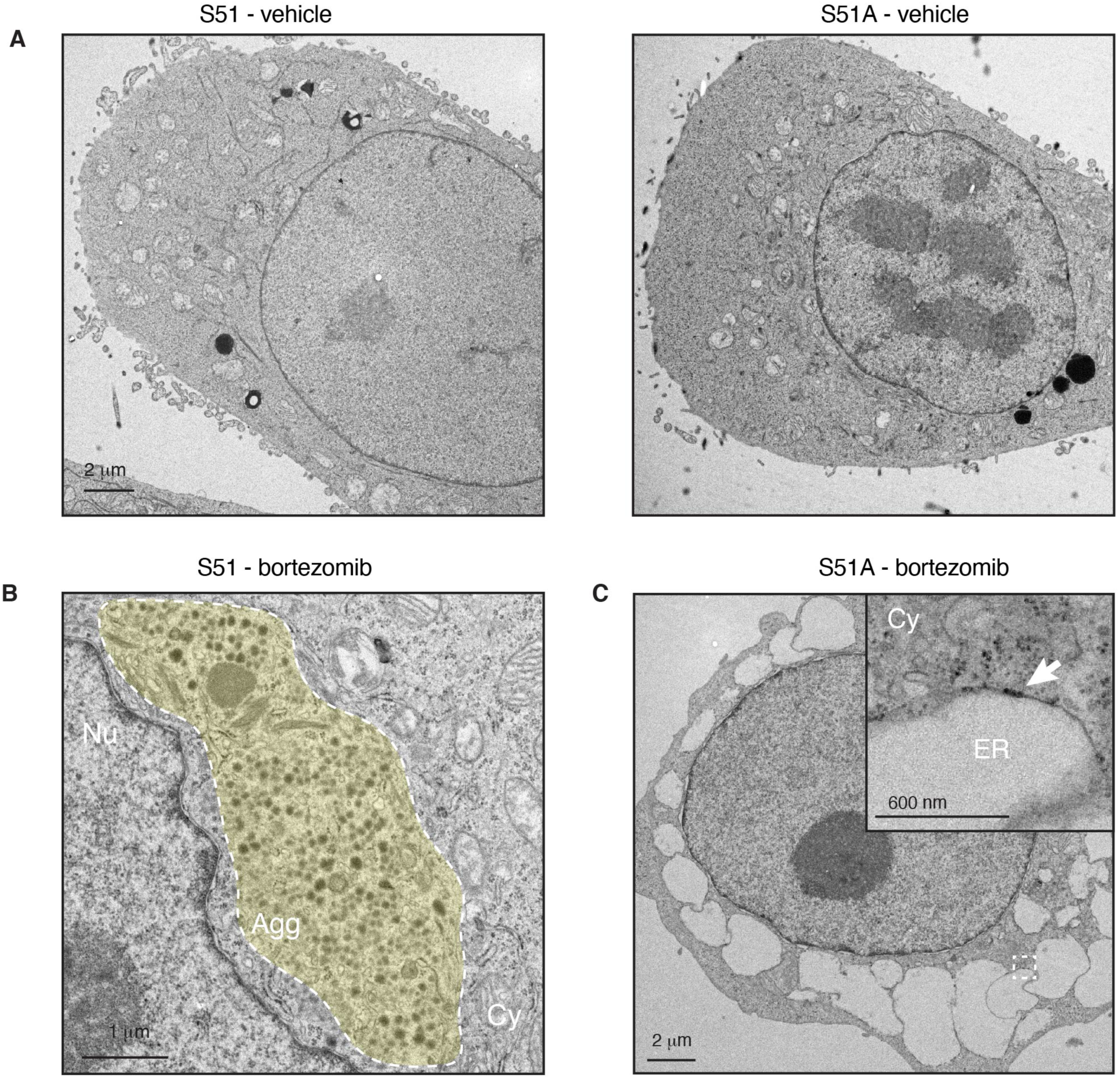
ISR promotes aggresomes formation upon recovery from proteotoxic stress. A. Transmission electron microscopy shows no difference at steady state between ISR-competent and ISR-null cells. No aggresomes are observed in the absence of bortezomib. Scale bar 2 μm. B. Transmission electron microscopy shows aggresomes, which form in S51 cells during recovery from proteotoxic stress. Aggresomes are highlighted by a yellow shadow. As expected aggresomes are perinuclear structures distorting the nucleus. Nu, nucleus. Cy, cytoplasm. Agg, aggresome. Scale bar 1 μm. C. Transmission electron microscopy reveals swollen ER that is present in some of the ISR-null cells following proteotoxic stress. These cells are viable and do recover from stress, but their presence is indicative of ER stress and unfolded protein that accumulate in the ER as well as the cytoplasm. White arrows indicate ribosomes attached to the ER membranes. ER, endoplasmic reticulum. Cy, cytoplasm Scale bar 2 μm.

**Figure 5-Figure supplement 2:**
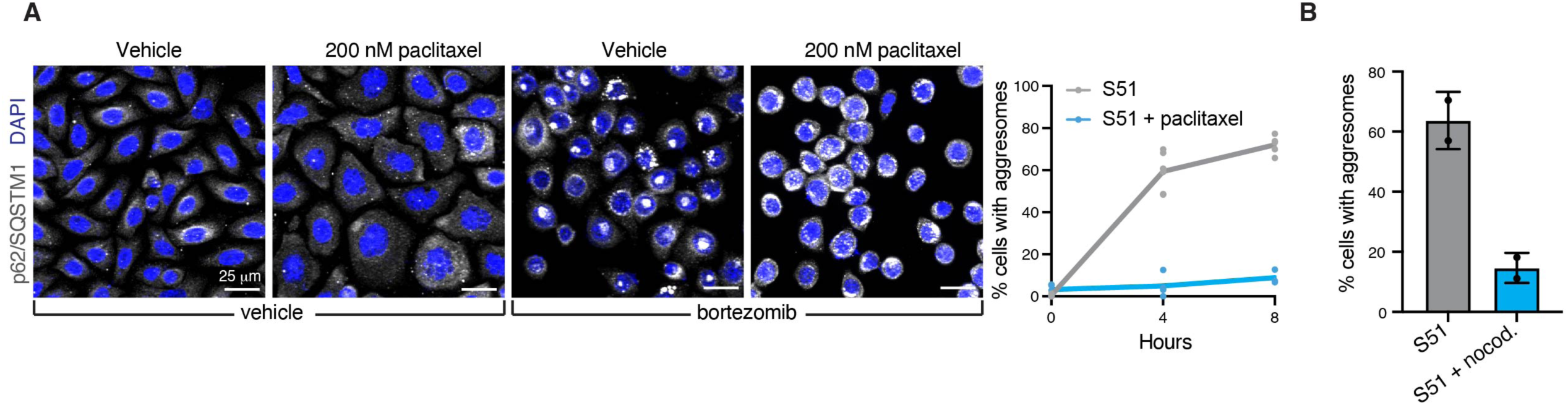
Aggresome formation is dependent on microtubule dynamics. A. Formation of aggresomes is dependent on microtubule-mediated transport. Blocking microtubule dynamics by using 200 nM of paclitaxel inhibits aggresome formation. Graph shows percentage of cells with aggresomes as quantified by immunofluorescence in two independent biological experiments. B. Aggresomes formation is dependent on microtubule-mediated transport. Microtubule polymerization was blocked adding 13 μM nocodazole at the time of bortezomib wash-out. Percentage of cells with aggresomes ± SD is quantified in two independent biological replicates.

**Figure 6-Figure supplement 1:**
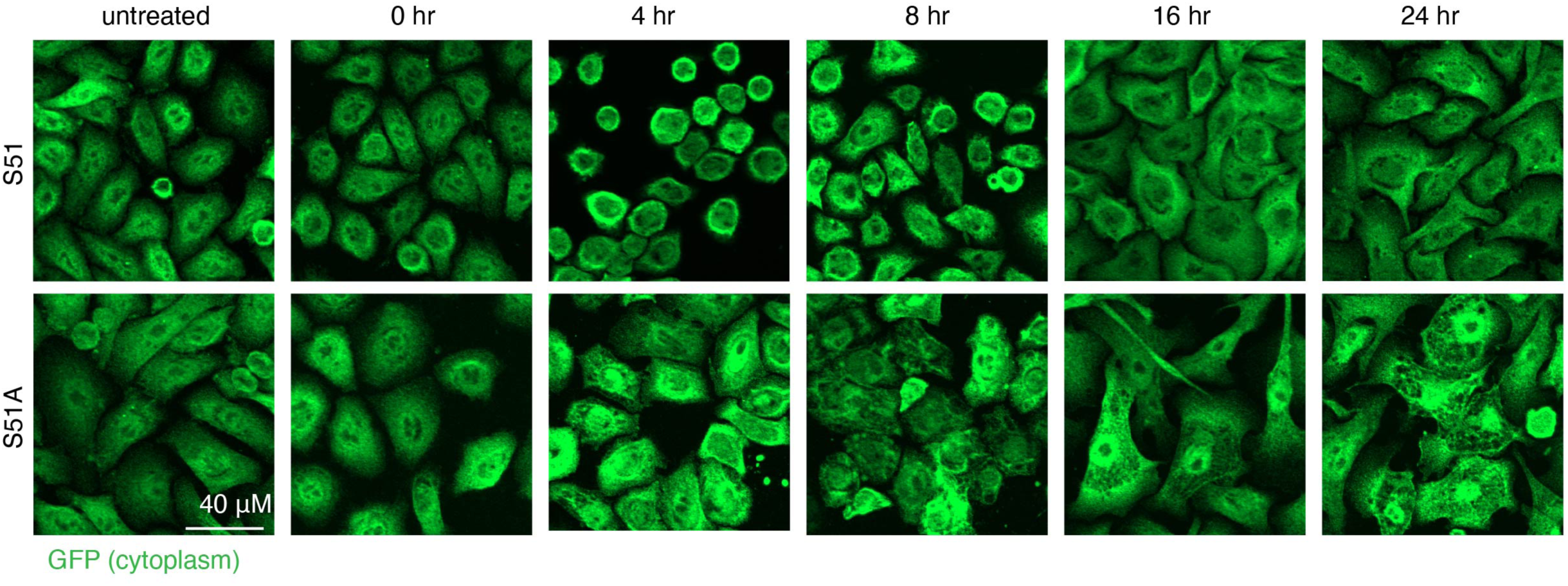
ISR promotes dramatic morphological changes in response to proteotoxic stress. ISR-competent or ISR-null cells are visualized via immunofluorescence by anti-GFP staining. Stress response in ISR-competent cells *in vitro* is characterized by a dramatic transient change in cell morphology, which is not observed in ISR-null cells.

## Source data

Figure 1C – Source Data 1

Raw immunoblots show eIF2*α* phosphorylation status in ISR-competent and ISR-null cell lines.

Figure 1D – Source Data 2

Raw immunoblots show protein synthesis rate in ISR-competent and ISR-null SCC cells upon indicated stresses.

Figure 1 – Figure supplement 1A – Source Data 3

Raw immunoblots show phosphorylation of eIF2*α* upon indicated stresses.

Figure 2E – Source Data 4

Raw immunoblots show ubiquitin and K48-linked polyubiquitin levels in ISR-competent and ISR-null SCC cells upon recovery from proteotoxic stress.

Figure 2 – Figure supplement 1B - Source Data 5

Raw immunoblots show eIF2*α* phosphorylation status of ISR-competent and ISR-null cells isolated from allografted tumors at day 35.

Figure 2 – Figure supplement 3A - Source Data 6

Raw immunoblots show ubiquitin levels upon recovery from proteotoxic stress in ISR-competent cells treated with the translation inhibitors cycloheximide or harringtonine.

Figure 2 – Figure supplement 3B - Source Data 7

Raw immunoblots show ubiquitin levels upon recovery from proteotoxic stress in ISR-competent cells treated with the autophagy inhibitor bafilomycinA1

Figure 3I – Source Data 8

Raw immunoblots show ATF5 and ATF4 levels in ISR-competent and ISR-null cells treated for 6 hrs with either vehicle or bortezomib.

Figure 7C – Source Data 9

Raw immunoblots show GFP-tagged ATF5 overexpression S51A-ATF5 cells.

Figure 7C – Source Data 10

Raw immunoblots show ubiquitin levels in the insoluble fraction of cells with indicated genotypes either vehcle treated of treated for 6hrs with bortezomib and let to recover for 24 hrs.

Source Data 11

Numerical data for each experiment included in the manuscript.

Source Data 12

Read counts for ribosome profiling and total RNA sequencing.

